# Optimal HLA imputation of admixed population with dimension reduction

**DOI:** 10.1101/2023.04.12.536582

**Authors:** Venceslas Douillard, Nayane dos Santos Brito Silva, Sonia Bourguiba-Hachemi, Michel S. Naslavsky, Marilia O. Scliar, Yeda A. O. Duarte, Mayana Zatz, Maria Rita Passos-Bueno, Sophie Limou, Pierre-Antoine Gourraud, Élise Launay, Erick C. Castelli, Nicolas Vince, SNP-HLA Reference Consortium (SHLARC)

**Affiliations:** Nantes Université, INSERM, Ecole Centrale Nantes, Center for Research in Transplantation and Translational Immunology, UMR 1064, F-44000 Nantes, France; São Paulo State University, Molecular Genetics and Bioinformatics Laboratory, School of Medicine, Botucatu, State of São Paulo, Brazil; Human Genome and Stem Cell Research Center, University of São Paulo, São Paulo, SP Brazil; Department of Genetics and Evolutionary Biology, Biosciences Institute, University of São Paulo, São Paulo, SP Brazil; Hospital Israelita Albert Einstein, São Paulo, SP Brazil; Medical-Surgical Nursing Department, School of Nursing, University of São Paulo, São Paulo, SP Brazil; Epidemiology Department, Public Health School, University of São Paulo, São Paulo, SP Brazil; Department of Pediatrics and Pediatric Emergency, Hôpital Femme Enfant Adolescent, CHU de Nantes, Nantes, France

**Author notes:** Correspondence, Nicolas Vince Nantes Université, CR2TI UMR1064 – ITUN, CHU Nantes Hôtel Dieu, 30 bld Jean Monnet, 44093, Nantes Cedex 01, France.

## Abstract

Human genomics has quickly evolved, powering genome-wide association studies (GWASs). SNP-based GWASs cannot capture the intense polymorphism of *HLA* genes, highly associated with disease susceptibility. There are methods to statistically impute *HLA* genotypes from SNP-genotypes data, but lack of diversity in reference panels hinders their performance. We evaluated the accuracy of the 1,000 Genomes data as a reference panel for imputing HLA from admixed individuals of African and European ancestries, focusing on (a) the full dataset, (b) 10 replications from 6 populations, (c) 19 conditions for the custom reference panels. The full dataset outperformed smaller models, with a good F1-score of 0.66 for *HLA-B*. However, custom models outperformed the multiethnic or population models of similar size (F1-scores up to 0.53, against up to 0.42). We demonstrated the importance of using genetically specific models for imputing admixed populations, which are currently underrepresented in public datasets, opening the door to HLA imputation for every genetic population.

## Introduction

Genome-wide association studies (GWASs) have now become a strong ally in the understanding of the underlying mechanisms of diseases susceptibility and outcomes, with historical associations such as rs2395029 in HIV (Limou and Zagury 2013; Fellay et al. 2007), or the identification of 233 genomic regions linked to multiple sclerosis susceptibility (International Multiple Sclerosis Genetics Consortium 2019). GWASs have also been performed as first lines of research at the beginning of the SARS-CoV-2 outbreak to evaluate how host genetics can influence COVID-19 outcomes (Pairo-Castineira et al. 2021; COVID-19 Host Genetics Initiative 2021; Douillard et al. 2021b; Castelli et al. 2022). Starting from the first GWAS with hundreds of individuals in the 2000s (Klein et al. 2005; Duerr et al. 2006), multiple initiatives emerged in the last decade seeking to systematically gather clinical and genetic information, such as the UK Biobank (Bycroft et al. 2018), Japanese BioBank (Hirata et al. 2017), or TOPMed (Taliun et al. 2021), which count hundreds of thousands of samples. These studies greatly improved the comprehension of the genetic impact on phenotype variation (Visscher et al. 2017; Tam et al. 2019; Claussnitzer et al. 2020). Along with the collective organization effort, continuous advances in the domain of Single Nucleotide Polymorphism (SNP) imputation (Browning et al. 2018), and the availability of computing power from imputation servers, globally helped the genomics community (McCarthy et al. 2016).

A bystander effect of these GWASs has been confirming the central role of the Major Histocompatibility Complex (MHC), especially the *HLA* genes, in immune-related diseases. The MHC was discovered in the 1950s (Dausset 1958), and was identified as crucial for transplantation success (Dausset 1981). Association studies expanded our understanding of the role of MHC since 2.5% of all significant associations in the GWAS catalog (MacArthur et al. 2017) coincide with the MHC region, and approximately 20% of all traits are associated with at least one SNP within the MHC (Douillard et al. 2021b). The associations go from auto-immune diseases such as type 1 diabetes (Concannon et al. 2009) or multiple sclerosis (International Multiple Sclerosis Genetics Consortium 2019), neurological disorders such as Parkinson(Nalls et al. 2019), to infectious diseases such as HIV (Limou et al. 2009; Fellay et al. 2007), Hepatitis B (Hu et al. 2013) and C (Vergara et al. 2019).

In the context of genetic association studies, a parallel effort focused on direct association with *HLA* polymorphisms to understand the mechanisms in which HLA molecules impact disease susceptibility and severity. These studies have identified protective and risk *HLA* alleles, such as *HLA-DRB1*15:01* in multiple sclerosis (Moutsianas et al. 2015), *HLA-DRB1*09:01* with tIgE levels in asthma (Vince et al. 2020b), specific HLA-DQB1 amino acids in hepatitis C virus infection (Valencia et al. 2022), or the HLA- DRB1 valine 11 in Parkinson’s disease (Domenighetti et al. 2022), among many others. The five most polymorphic *HLA* genes (*HLA-A*, *HLA-B*, *HLA-C*, *HLA-DQB1,* and *HLA-DRB1*) are exceptionally diverse, with almost 30,000 alleles combined (Robinson et al. 2020). However, most of these alleles seem to have frequencies <1% (Maiers et al. 2007). Therefore, because of the high number of alleles and their low frequency, the HLA typing of thousands of individuals is necessary to reach sufficient statistical power for detecting associations. The cost-efficiency of directly typing *HLA* for such cohorts is limited. Thus, following the steps of the SNP association, the HLA community organized multiple typing initiatives and developed imputation tools (Meyer and Nunes 2017; Douillard et al. 2021a). The literature on HLA imputation articulates a dual focus on algorithms and reference data.

Regarding algorithms, several *HLA* imputation tools allow to create reference panels for imputing *HLA* alleles from SNP data: HIBAG (Zheng et al. 2014) and SNP2HLA (Jia et al. 2013) are the most common choices. Pappas et al. evaluated HIBAG to be the most accurate (Pappas et al. 2015). A new generation of software followed, with improvements to existing algorithms such as HLA*IMP:03 (Motyer et al. 2016) and CookHLA (Cook et al. 2021), or using deep learning with DEEP-HLA (Naito et al. 2021), all of which will probably gain traction over time. However, regarding reference datasets, the accuracy of *HLA* imputation results depends on the reference panel used to predict the target genotypes; if training and target data are not of the same ancestry, it will provide inaccurate results due to different HLA alleles and linkage disequilibrium patterns between SNP and HLA in different populations. To circumvent this issue, researchers advocated for both: specific reference panels, such as in Japan (Okada et al. 2015), Finland (Ritari et al. 2020), or SweHLA (Nordin et al. 2020), and large multi-ethnic reference panels (Degenhardt et al. 2019; Luo et al. 2021). To pursue the different efforts, we created the SNP-HLA Reference Consortium, or SHLARC (Vince et al. 2020a). Our goal is to coordinate an international effort to gather HLA data and reference panels, make them available to the scientific community and improve the methodology of *HLA* association studies. Generally, *HLA* imputation is highly performant for European-origin populations as a large amount of data are available to build reference panels. Conversely, the challenge is higher when focusing on admixed or underrepresented populations as fewer data are available. A clear HLA imputation strategy remain to be defined to improve accuracy in these populations: here, we want to increase our understanding about HLA imputation performance between larger reference panels or smaller but customized (ancestry-matched) reference panels. Indeed, our hypothesis is that oversampling individuals for the reference panel with close genetic ancestry to the target individuals would increase accuracy for their specific *HLA* alleles. To explore this in our study, we focused on the results of *HLA* imputation on admixed populations using a multiethnic reference panel from the 1,000 Genomes Project (1KG), and investigating dimension reduction as a method to mitigate *HLA* imputation errors on rare alleles (1000 Genomes Project Consortium et al. 2015; Byrska-Bishop et al. 2022).

## Results

### HLA imputation strategy

HLA imputation accuracy heavily depends on the data used as reference. Our study aims at finding the preferred *HLA* imputation combination of reference data selection and imputation method when dealing with a target population whose ancestry is absent or underrepresented in the available training data. The 1KG dataset presents a large diversity in populations as described in table S1 (1000 Genomes Project Consortium et al. 2015; Clarke et al. 2017), which can be grouped in 5 populations: African (AFR), American (AMR), European (EUR), East Asian (EAS), and South Asian (SAS). We selected these data as a training dataset to create 395 reference panels to be tested (Figure 1), including: (a) the full dataset (full1KG, N=2,504), (b) 10 replications from 6 populations (1KG, AFR, AMR, EUR, EAS, and SAS; N=200 for each), (c) 19 conditions for the custom reference panels (further described in the next chapter; 200<N<485); each condition replicated 5 times for each *HLA* gene (*HLA-A*, *HLA-B*, *HLA-C*, *HLA-*DQB1, and HLA-DRB1).

**Figure 1.**
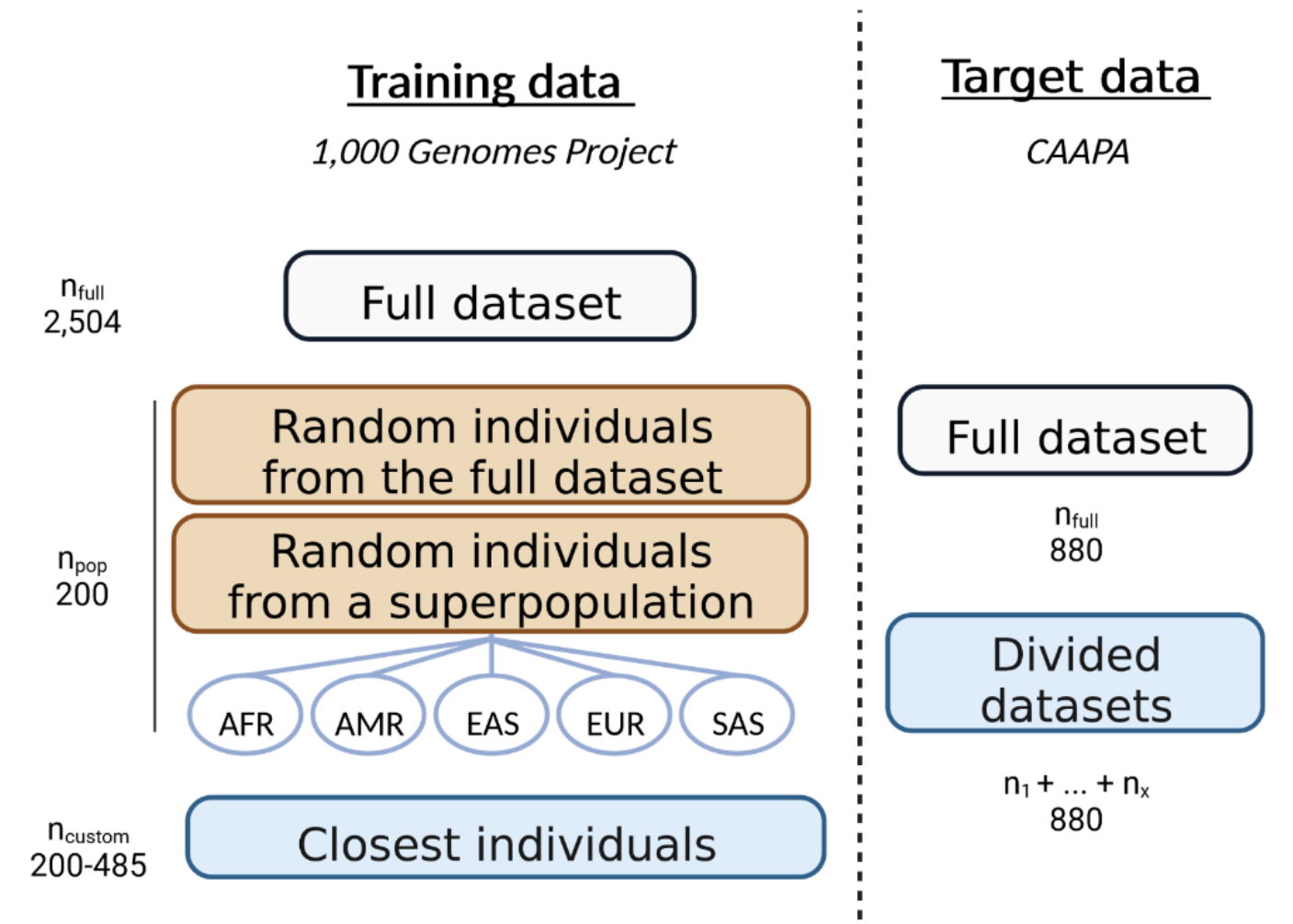
Selection strategy: description of the dataset selection for training and testing. Different subsets of the 1KG dataset are used as reference, selected by super-population or from genetic proximity with the CAAPA dataset. The HLA genotypes from CAAPA are either imputed from a single model or by multiple models specific to subsets of CAAPA.

The CAAPA cohort (Consortium on Asthma among African-ancestry Populations in the Americas) is constituted of 880 individuals with SNPs of the *MHC* region and HLA genotypes. These individuals are from admixed African and European ancestry in various proportions (Vince et al. 2020b). Only a small fraction of these populations ancestries are represented in the 1KG dataset, so we also wanted to evaluate the impact of admixture in the imputation process and accuracy. Thus, the CAAPA population was alternatively considered a unique dataset of 880 individuals, or as multiple subsets of it, depending on the representation with dimension reduction methods.

#### Data selection for customized HLA imputation

We created models with individuals from 1KG genetically close to the CAAPA target data: the custom models. We decided to rely on dimension reduction, common in population genetics, to assess individuals ancestry. The goal is to select 200 individuals from 1KG closest to the target data, regardless of their designated ancestry. Classically, ancestry is assessed with whole-genome SNPs by Principal Component Analysis (PCA). However, since we focused our study on HLA, we decided to represent the populations using only SNPs within the MHC region (29-34Mb from chr6). This representation strategy separated the African population and a portion of the American population in one part, and the rest of 1KG on the other (Figure 2A). The usual granularity of PCA on whole-genome genotypes (Figure S1A) is not obtained and does not allow grouping ancestries. However, we could identify well-separated groups with a two-dimension UMAP (Uniform Manifold Approximation and Projection) of the MHC region (Figure 2B; UMAP on whole-genome genotypes in Figure S1B).

**Figure 2.**
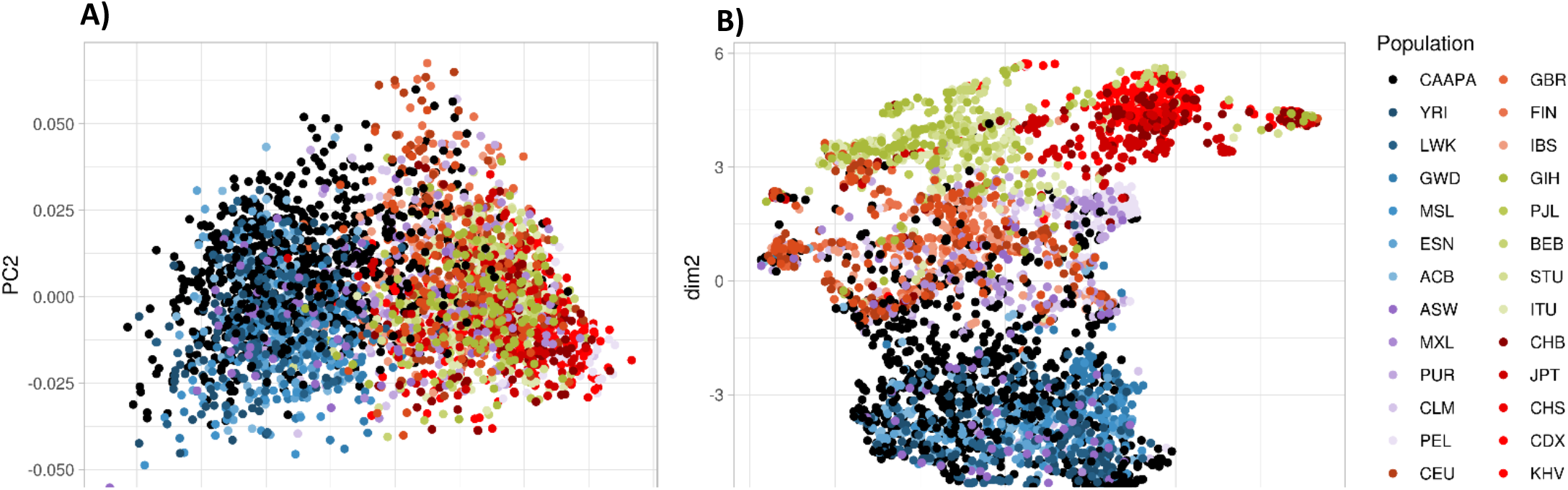
PCA (A) and UMAP (B) representation of 1KG and CAAPA dataset with merged genotypes of the MHC region. CAAPA is represented in black. Super-populations are colored in five main colors divided into different shades for each population (Table S1), including 5 super-populations: African (AFR) in blue, American (AMR) in purple, European (EUR) in orange, South Asian (SAS) in green, East Asian (EAS) in red. PCA does not separate well the population when restrained to the MHC region, whereas UMAP creates different groups of ancestries.

To investigate the effect of dimension reduction on HLA imputation, we tested 3 parameters for representation: the algorithm (PCA or UMAP), the number of dimensions used (2 or 10), and the genomic region covered by the genotypes dataset (the whole chromosome 6 or the MHC region, see also Figures S1). The different conditions are named after the combination of these parameters. For instance, a selection of the training data based on a UMAP using the distance computed in 10 dimensions on every SNP available on chromosome 6 is named UMAPnonMHC_10D.

We performed a silhouette score analysis to the resulting projection of the CAAPA dataset. We identified that, in every UMAP condition and with the ten-dimensions PCA in the MHC region, we could cluster CAAPA in more than one group. In these cases, we decided to create one model per group. We computed the average coordinates of the CAAPA individuals, then selected the 200 individuals from 1KG closest to this point (Figure 3). To avoid redundant models, we checked the overlap of selected individuals between the conditions. Surprisingly, they all yielded a unique list of 1KG individuals, with low overlap between conditions (Figure S2). For the conditions where the CAAPA dataset was separated into different subsets, we imputed the individuals separately, thus relying on multiple models, but merged the results into one table. For example, with the two-dimension UMAP representation (genotypes from the MHC region), we computed three different models of 200 1KG individuals (Figure 3). We then imputed three CAAPA groups independently (357, 344, and 179 individuals for a total of 880) and combined results into a unique table of imputation for the full dataset.

**Figure 3.**
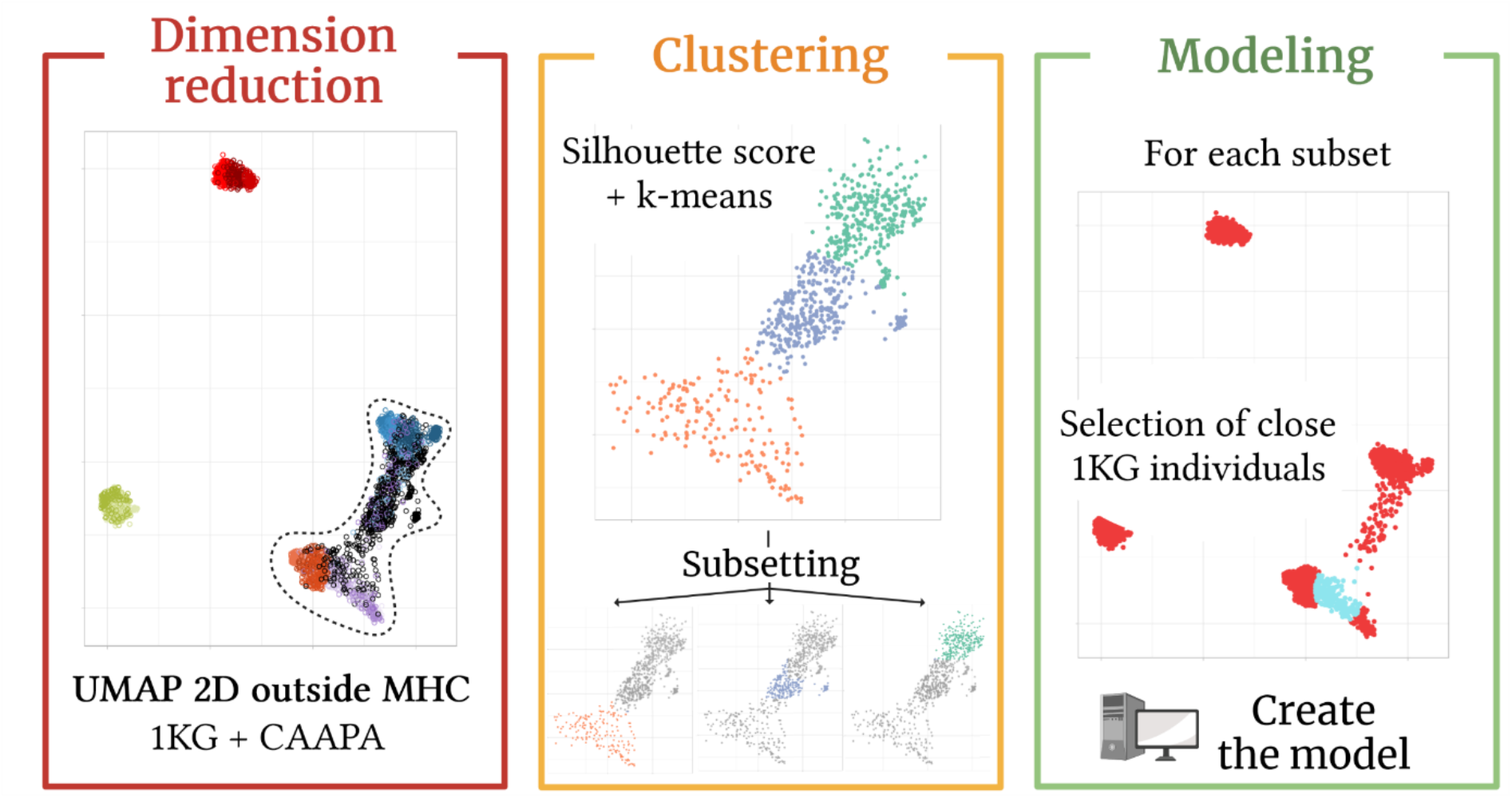
Creation of custom 1KG models for CAAPA imputation. 1) <u>Dimension reduction</u> allows the separation of individuals according to ancestry, using UMAP as an example. CAAPA is represented in black. Super-populations from 1KG are colored in five main colors: African (AFR) in blue, American (AMR) in purple, European (EUR) in orange, South Asian (SAS) in green, East Asian (EAS) in red. It is also possible to apply dimension reduction to one dataset and project another onto it. 2) <u>Clustering</u> of the target dataset: here CAAPA. The silhouette score allows to evaluate the preferred number of clusters, then k-means allows for subsetting. 3) <u>Modeling</u>. The barycenter of each cluster subset is computed, then 1KG individuals closest to this coordinate are selected (in light blue), allowing to create a custom model. Created with biorender.com

### CAAPA *HLA* imputation comparison between usual and custom reference panels from 1KG data

We have compared the different conditions by averaging the F1-score of each allele. As explained by Cook et al. (Cook et al. 2021), the F1-score has an advantage over other accuracy metrics for representing the rare alleles as it is the mean of two metrics, taking into account both the potential under- and over-prediction of an allele. As expected, the full1KG model (N=2,504) displayed the highest F1-score for all *HLA* genes, ranging from 0.64 for *HLA-DRB1* to 0.87 for *HLA-C* (Figure 4). For *HLA-B*, full1KG has a score of 0.66. However, still for *HLA-B*, when considering the smaller models, we found that the 1KG models (F1-score of 0.42) and the populations with close ancestry to CAAPA (AFR: 0.37, AMR: 0.42) had nominally lower F1-score than some custom models (PCAnonMHC_10D: 0.52, UMAPnonMHC_10D: 0.53). This trend was also observed for *HLA-A* and *HLA-DRB1*, while *HLA-C* and *HLA-DQB1* show a higher mean F1-score for the small 1KG models. F1-scores are not to be interpreted as regular accuracies. Indeed, when the same methodology is applied to compute the average accuracies of each allele, these accuracies obtained more than 98% (represented as error rates in Figure S3). Additionally, the individual and haplotype accuracies, which corresponds to the proportion of correct genotypes (individuals can be counted as 0 or 1; incorrect vs. correct imputation) and the proportion of correct allele (individuals can be counted as 0, 0.5, or 1; incorrect vs. 1 correct allele vs. 2 correct alleles imputation), respectively, also show values above 80% (Figure S4).

**Figure 4.**
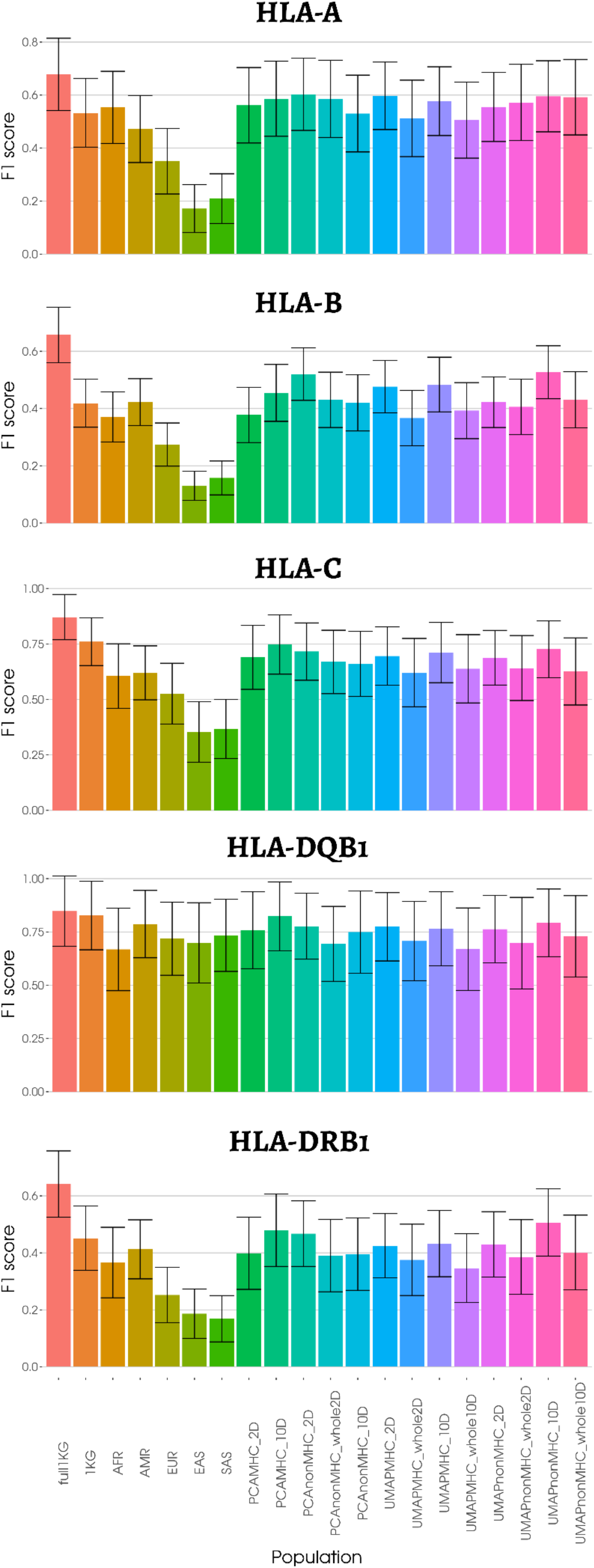
Average F1-score of HLA allele predictions for HLA-A, HLA-B, HLA-C, HLA-DQB1, and HLA-DRB1 based on imputation of the CAAPA dataset, with different training models from the 1000 Genomes dataset. We have removed alleles that are not represented in the training datasets. Nomenclature of the models can be found in *table S2*. Full1KG, N=2,504. Small 1KG models (1KG, AFR, AMR, EUR, EAS, SAS), N=200. Custom models, 200<N<485.

To investigate the impact of custom models on imputation and why they seemed to perform better for highly polymorphic genes, we stratified the mean F1-score metric by *HLA* allele frequency (Figure 5). The full1KG model (N=2,504) yielded a higher F1-score through all allelelic frequencies. Custom models performed equally or marginally better for the rarest alleles (frequency <= 0.1%) and the most common alleles (frequency >10%). Still, they scored higher for every other category than population models. For *HLA-B*, UMAPnonMHC_10D (N=485) presented an F1-score of 0.30, 0.70, 0.85, and 0.91 for the categories from 0.1 to 10% frequency, whereas the multi-ethnic model (1KG, N=200) showed scores of 0.18, 0.45, 0.78, and 0.91. Notably, the reference panel based on the African population performed worse for *HLA-DQB1.* It can be explained by the allele *HLA-DQB1*06:01*, which was represented only once and had an F1-score of 0.1.

**Figure 5.**
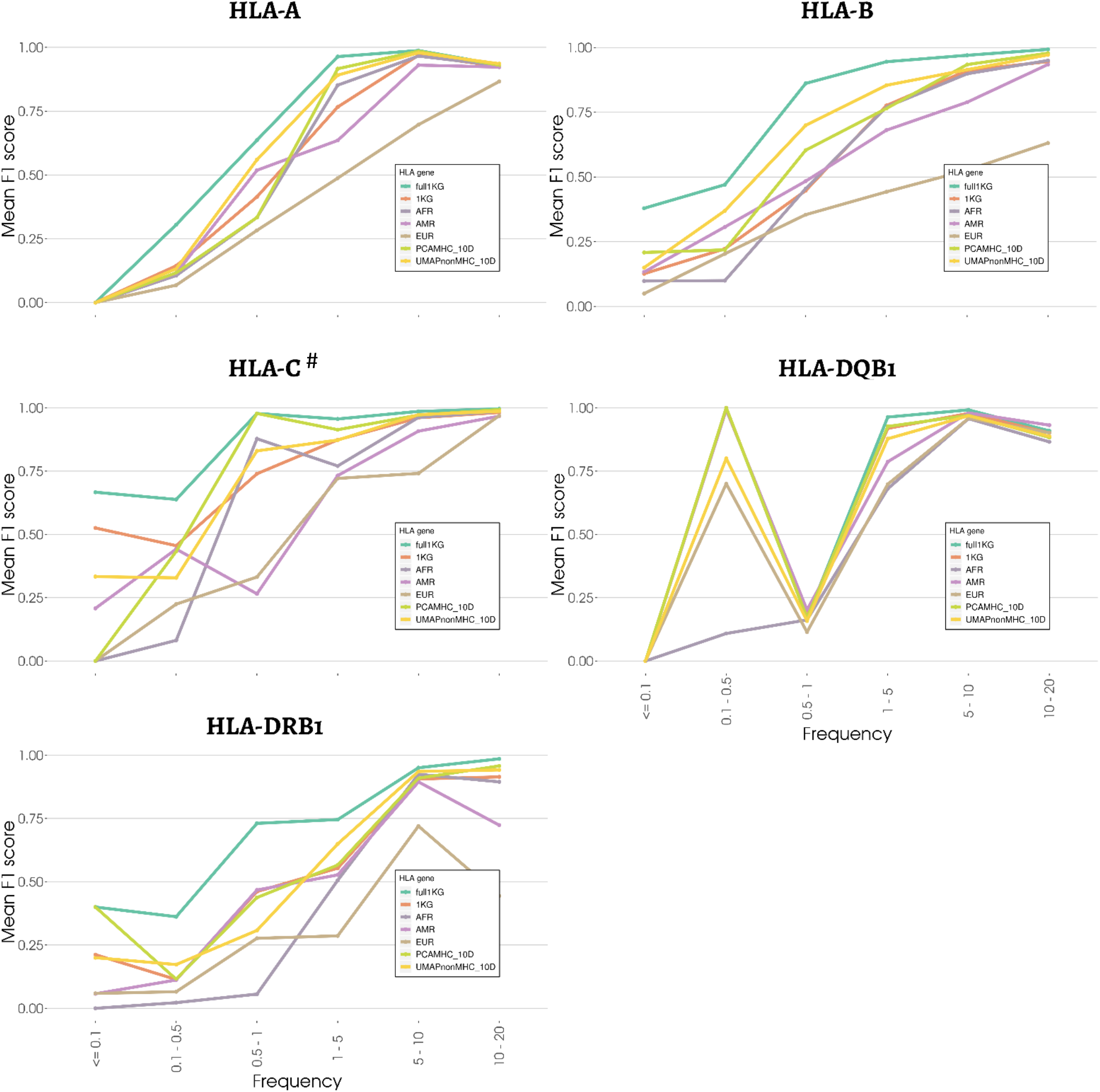
Mean F1-score of HLA alleles imputation, stratified by groups of frequency, for the full 1KG model, super- population models, and a selection of custom models. The custom models PCAMHC_10D and UMAPnonMHC_10D are displayed as they are the most accurate. #: HLA-C does not have the 10-20% frequency category. Therefore, unlike other genes, the last two categories correspond to 5-10% and >20%.

The results showed that creating custom reference panels based on a genotypic distance between individuals can improve the outcome compared to multi-ethnic or declared ancestry panels. However, larger multi-ethnic reference panels are always more robust. We went further and looked directly at the imputation of *HLA* alleles individually.

When we analyzed results allele by allele, taking *HLA-A* (Figure 6) as an example, we observed that in most cases, custom models performed just as well, or a few points under the full dataset models (e.g. *HLA-A*01:01*, *HLA-A*23:01*). Several *HLA* alleles were better predicted with the custom models compared to the multi-ethnic (1KG) and population models (e.g. *HLA-A\**01:02, *HLA-A\**80:01), highlighting the importance of creating specific reference panels. We found cases where the full1KG model (N=2,504) or population models (N=200) were the only ones to predict the allele (e.g. *HLA- A\**02:06, *HLA-A\**03:02). However, we also found cases where custom models were the only ones to impute correctly the allele (e.g. *HLA-A*02:04*). Zheng et al. (Zheng et al. 2014) showed that at least 10 copies of an allele were needed in a model to be able to impute them. Nine *HLA-A* alleles were present in the training and target data but were not imputed by any of the models (e.g. *HLA-A*02:11*, *HLA- A*24:03*, *HLA-A*26:08*). Often, the allele was present only in few individuals of the target data, causing the miscalled allele to weigh a lot in the score. We focused on *HLA-A* for visualization purposes, but the results applied to *HLA-B*, *HLA-C*, and *HLA-DRB1* (Figure S5). Interestingly for *HLA-DQB1*, the best custom models were never the best predictors. For most *HLA-DQB1* alleles, the best training dataset was the full1KG dataset. For *HLA-DQB1*03:01,* however, the AMR and EUR super-populations yielded better results. The different examples presented here show how a custom reference panel could help in the imputation of certain *HLA* alleles. However, since bigger models produce better imputation results overall, we would need to know when to select the results from the custom reference panel.

**Figure 6.**
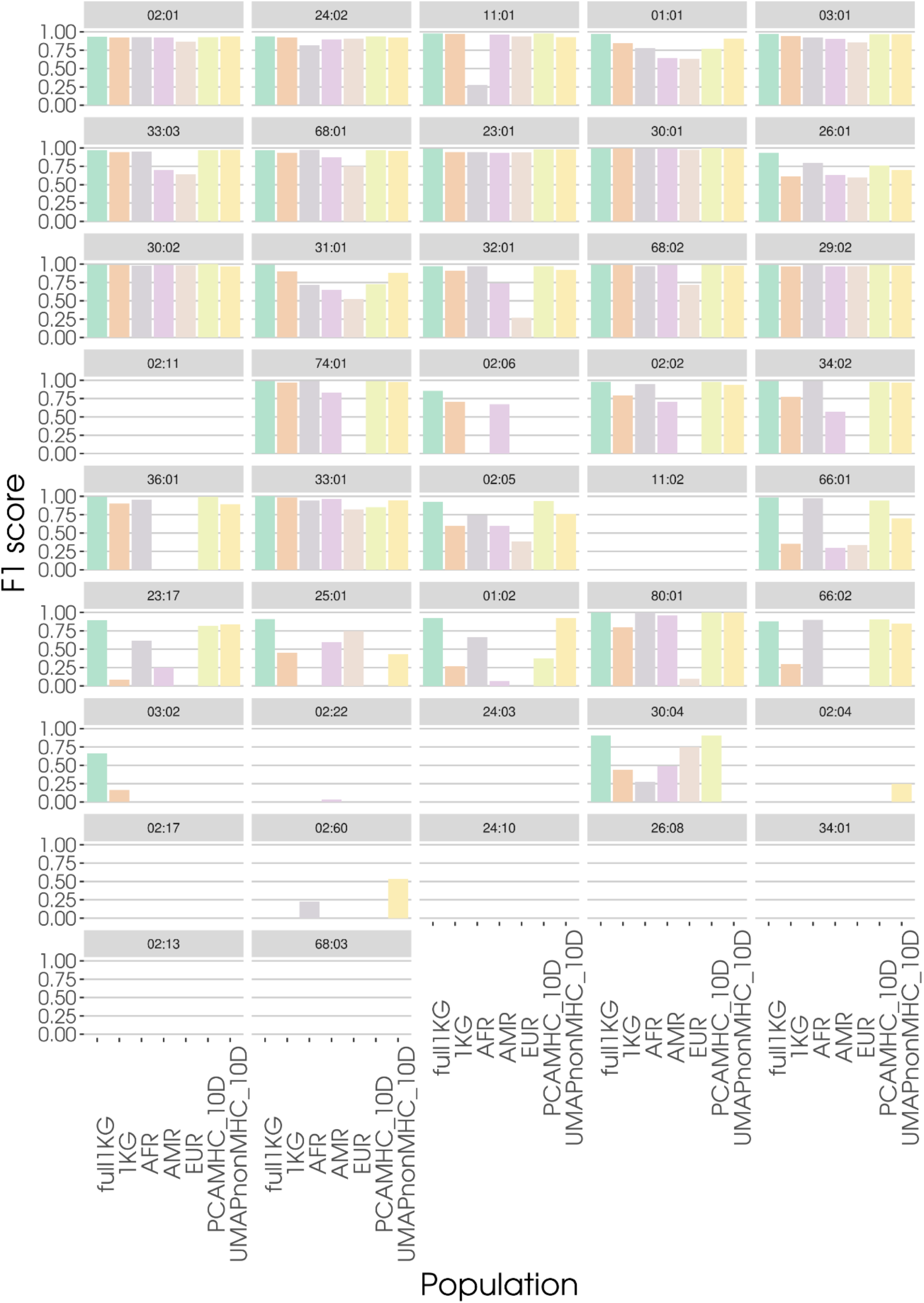
Mean F1-score of each HLA-A alleles (N=42) for the full 1KG training dataset, the African, American, European super-populations datasets, and the most accurate custom reference panels PCAMHC_10D and UMAPnonMHC_10D. Alleles are ordered by decreasing frequency in the 1KG dataset. Those absent from the training dataset have been removed to compute the means.

HLA imputation with HIBAG yields post-probabilities for each genotype. We tried to harness the few cases where custom models performed better (in terms of post-probabilities) to obtain hybrid imputation between the full models and the custom model. We chose UMAPnonMHC_10D as it performed the best on multiple *HLA* genes. Unfortunately, the small number of samples in the custom models led to lower post-probabilities than the full model. In the few cases where UMAPnonMHC_10D yielded better post-probabilities, the imputed genotype was not always correct, whereas the less likely genotype imputed by the other model was correct. In a real situation where the *HLA* alleles of the target data would not be known, there would be no way to choose between the imputed genotype of the two models (Figure S6).

### Replication with admixed Brazilian individuals from SABE

We replicated our methodology on another cohort of admixed individuals, the Longitudinal Health, Well-Being, and Aging cohort (SABE - *Saúde, Bem-estar e Envelhecimento*) from Brazil, to validate the impact of the models composition on *HLA* imputation (Figure 7). SABE is an independent dataset of 1,322 individuals from Brazil, mostly with European and African admixed ancestry (Naslavsky et al. 2022). To validate our conclusions, we used the same models as with the CAAPA dataset; therefore, between 11.6% and 45.1% of the model SNPs were missing in the target data. Though it probably reduced the imputation score overall, the missing SNPs were homogeneous across conditions for each gene, with averages of 30,0% for *HLA-A*, 14,3% for *HLA-B*, 13,9% for *HLA-C*, 39,4% for *HLA-DQB1*, and 39,6% for *HLA-DRB1*. We also limited our study to the PCAMHC_10D and UMAPnonMHC_10D custom models, as these two models predicted *HLA-A*, *HLA-C*, *HLA-DQB1,* and *HLA-DRB1* better, out of all the custom models in the CAAPA dataset.

**Figure 7.**
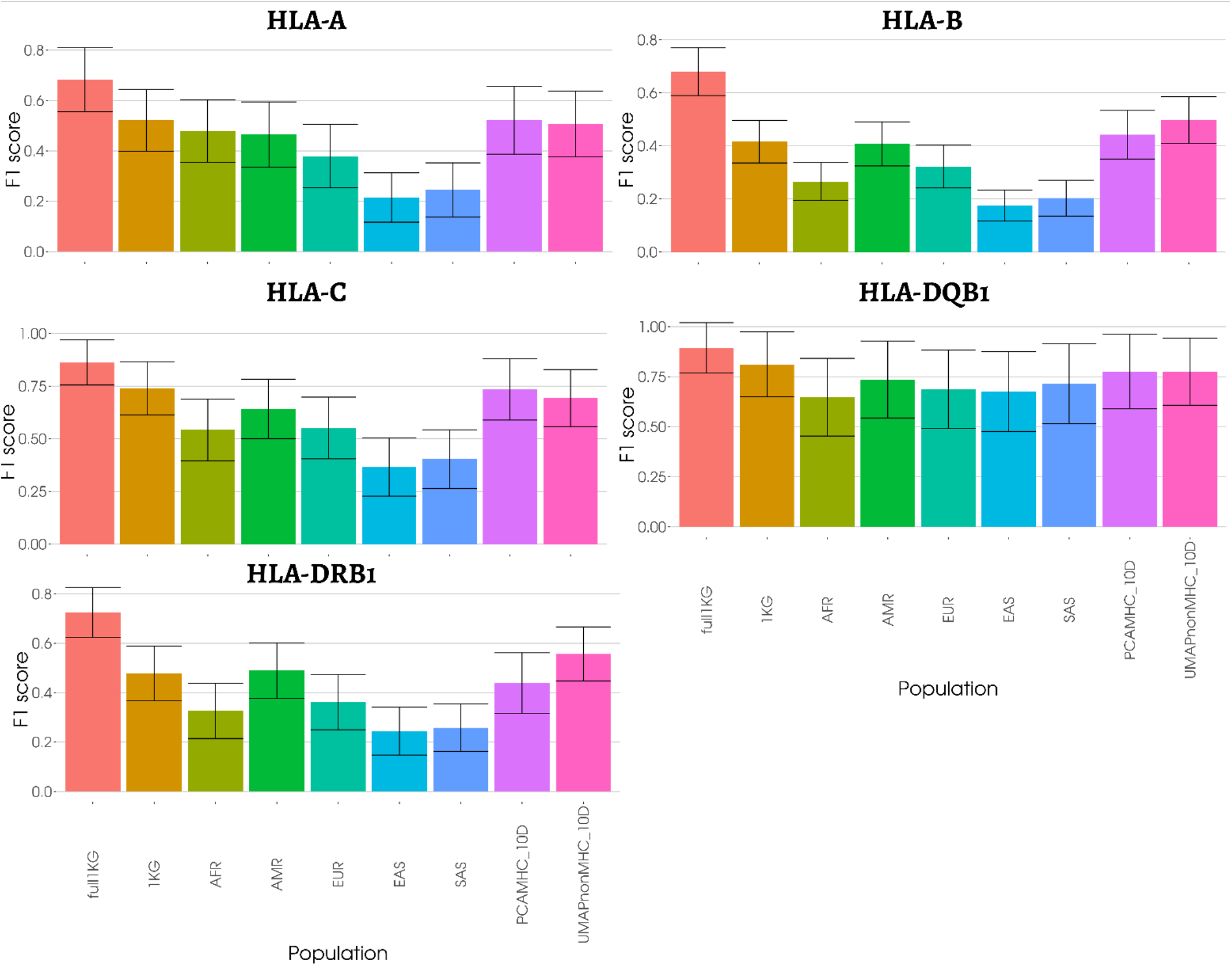
Mean F1-score of SABE’s imputed HLA-A, HLA-B, HLA-C, HLA-DQB1, and HLA-DRB1 genotypes, using the full 1KG model, compared to super-populations from 1KG or individual custom models selected by dimension reduction. Alleles absent from the training datasets were removed to obtain these values.

As with CAAPA, the custom models had nominally higher F1-score than the 1KG model, but only for the *HLA-B* (0.44, 0.50 for PCA and UMAP vs. 0.42 for 1KG) and *HLA-DRB1* (0.56 for UMAP vs. 0.48 for 1KG). Overall, the validation with the SABE population showed the same patterns as the CAAPA population, with a global preference for the full1KG model and multiple cases where the custom reference panels were to be preferred but presented low post-probabilities genotypes.

## Discussion

The HLA and immunogenetic community, along with the SHLARC (Vince et al. 2020a), carries a broad dynamic to provide scientists with reliable tools and reference panels for *HLA* imputation, thus increasing the power of *HLA* association studies to that of existing GWASs. We believe our results contribute to this effort. Our work focused on improving existing methods of *HLA* imputation by finely accounting for ancestry in the choice of the training model. Our underlying hypothesis was that oversampling individuals to create reference panels with close genetic ancestry compared to the target individuals would increase HLA imputation accuracy for rare *HLA* alleles. In this context, we chose to evaluate the imputation of CAAPA, an admixed African-American cohort, using reference panels composed of different combinations of 1,000 Genomes Project individuals: randomly selected from a population or selected for their estimated ancestry by dimension reduction. We showed that, ultimately, the number of individuals was the crucial point of HLA imputation. The reference panel composed of 2,504 individuals from 1KG systematically had a higher F1-score than other smaller models. Using fewer individuals for training by selecting individuals close to the ancestry of the target population was a good strategy and resulted in slightly better HLA imputation F1-scores, compared to multi-ethnic reference panels. The improvement did not concern the rarest or most common alleles, which are respectively badly and well imputed by all those models. At the allele level, we expected the full model to impute *HLA* alleles other models would not; we also saw the opposite with custom reference panels capturing a part of the information left out in the full model. Unfortunately, we could not conclude on its applicability since the custom reference panels had fewer individuals resulting in lower post-probabilities that rendered a hybrid imputation impossible. Research on SNP to SNP imputation also encounters the problem of lack of diversity for the imputation of rarer alleles, and are working with specific reference panels to enhance imputation accuracy (Kals et al. 2019; Herzig et al. 2022).

Interestingly, we were also able to use UMAP for genomic ancestry representation, as can also be seen in recent research (Diaz-Papkovich et al. 2021; Sakaue et al. 2020; Dai et al. 2020). It presented a good separation of ancestry groups in two dimensions when only using the MHC SNPs, concordant with the frequency difference of *HLA* alleles between populations (Maróstica et al. 2022). In contrast, PCA would fail to separate them in only two dimensions, limiting the possibility to visualize. PCA uses SNPs to explain most of the variance. Conversely, UMAP tries to preserve the topography of the higher dimensions in its reduction, taking into account every SNP available for distance. Besides, we observed a distance between individuals sometimes higher inside a labeled 1KG population than between populations, as described in Maróstica et al. (Lewontin 1972; Maróstica et al. 2022). This representation of this genomic diversity inside the MHC directly impacts how we should construct reference panels in the future and highlights the importance of gathering more data from different ancestry backgrounds.

Our work showed the potential interest of population-specific reference panels, as multiple studies have demonstrated (Okada et al. 2015; Ritari et al. 2020; Nordin et al. 2020; Luo et al. 2021; Mimori et al. 2019; Zhou et al. 2016; Nunes et al. 2016; Huang et al. 2020). However, we strayed further from the geographic definition of the population. We tried to find a local definition of ancestry to select training datasets. While doing so, we also omitted potential sides to the problem and created limits to our method. One important difference to *HLA* imputation compared to typing, inherent to the method, is the impossibility of predicting *de novo* alleles and the difficulty of imputing rare alleles. This issue is intrinsic to all training machine learning methods, and it is especially true for HLA, where each gene can have thousands of alleles. In HIBAG, for instance, an allele should be present at least 10 times in the training dataset to be predicted (Zheng et al. 2014). This study showed that this limit can be overcome to a certain extent but still hinders *HLA* imputation accuracy overall. Additionally, the choice to limit the number of randomly selected individuals was directly linked to the maximum of samples in the smallest population (n_AMR_=347). However, it has led to low imputation scores. Even though we performed replications, the difference between population models and the full dataset, or the custom models, may greatly vary if we increase this limit with another multi-ethnic dataset. It is one potential improvement to this work, which may validate or not our findings.

We chose to represent the *HLA* imputation with the F1-score, as seen in Cook et al. (Cook et al. 2021). This choice is convenient for the analysis of *HLA*, in which we encounter low and unbalanced frequencies between the different alleles. We set the F1-score at 0 when a specific allele was not imputed at all (whereas F1-score should be null) to represent all alleles in common between the two datasets and weigh this absence of imputation negatively. It has increased the confidence interval of each averaged F1-score and limited the possibility to find statistical differences between them. It is important to note that the F1-score gives a harsher view on *HLA* imputation because rare alleles have low scores, however, HLA imputation performs very well for common alleles (Figure S3) (Meyer and Nunes 2017).

Besides methodology, *HLA* imputation gains much accuracy from the number of samples and the diversity in the reference panels. This is why initiatives looking into expanding the *HLA* data and creating larger reference panels, such as Degenhardt et al., are essential to the field (Degenhardt et al. 2019; Luo et al. 2021; Abi-Rached et al. 2018). With the SHLARC (Vince et al. 2020a), we advocate for the coordination of such efforts to provide multi-ethnic panels of sufficient size, and help researchers do *HLA* imputation to investigate HLA risk and protection alleles, focusing on the coverage of the globe for data gathering. The evolution of imputation tools will also consequently improve *HLA* imputation. HLA-IMP*03 (Motyer et al. 2016) and CookHLA (Cook et al. 2021) showed improved results over the algorithms they are created upon, and DeepHLA (Naito et al. 2021) also showed high accuracy, with a specific focus on rare HLA alleles. Eventually, these efforts will reach a limit, and we think the main focus of research should be gathering data worldwide.

Our results demonstrated the interest of using genetically specific models for imputing admixed populations which are currently underrepresented, opening the door to *HLA* imputation for every genetic population, while also exemplifying some limitation. The SNP-HLA Reference Consortium (SHLARC) wants to contribute to the *HLA* association analysis community by providing a platform for *HLA* imputation with exhaustive and diverse reference panels. We hope this will help association studies to rapidly increase their statistical power and become a natural extension of genome-wide association studies pointing towards *HLA* association.

## Methods

### Data description and processing

SNPs data from the 1KG, CAAPA, and SABE cohorts were obtained from whole-genome sequencing. The 1KG dataset is one of the most diverse public dataset with 2,504 individuals from 26 populations (1000 Genomes Project Consortium et al. 2015; Clarke et al. 2017). These populations are grouped in 5 populations, as described in table S1: African (AFR), American (AMR), European (EUR), East Asian (EAS), and South Asian (SAS). *HLA* genotyping for the *HLA-A*, *HLA-B*, *HLA-C*, *HLA-DQB1,* and *HLA-DRB1* genes was published and made accessible using HLA calling algorithms for whole-genome sequencing data (Abi-Rached et al. 2018). Moreover, the SNP data has been updated with a new whole-genome sequencing of 30X coverage from the New York Genome Center (Byrska-Bishop et al. 2022). The CAAPA cohort (Consortium on Asthma among African-ancestry Populations in the Americas) was created to study asthma in African-ancestry populations. The aim of this study was to catalog genetic diversity in these populations, especially the African Diaspora in the Americas. From this, we had access to 880 individuals with whole-genome sequencing data of the *MHC* region and HLA genotypes (Vince et al. 2020b). The *HLA* alleles were called with the Omixon software (Budapest, Hungary) from whole-genome sequencing data (Vince et al. 2020b). The SABE (*Saúde, Bem-estar e Envelhecimento*) data come from the longitudinal, census-based follow-up, Health, Well-Being, and Aging cohort of elderly people from São Paulo, Brazil. SABE is an independent dataset of 1,322 admixed individuals from Brazil, mostly with European and African admixed ancestry: details can be found in the whole-genome sequencing flagship publication (Naslavsky et al. 2022). *HLA* genotypes for SABE cohort were obtained after read alignment with hla-mapper 4.1. This application was designed to optimize the mapping of *HLA* sequences produced by massively parallel sequencing procedures (Castelli et al. 2018); the pipeline is available at https://github.com/erickcastelli/HLA_genotyping/tree/main/version_2.

SNPs data were handled with PLINK v1.90b6.21 (Chang et al. 2015) and went through the same quality control step: the removal of A/T and G/C ambiguous SNPs, and SNPs with >2% missing genotypes and <1% minor allele frequency. *HLA* data comprises two-field alleles for *HLA-A*, *HLA-B*, *HLA-C*, *HLA-DQB1,* and *HLA-DRB1*, stored in a CSV file. *HLA* imputation models were computed on R 3.5.3 (R Core Team 2022) with HIBAG v1.19.3 (Zheng et al. 2014) and its complementary package HIBAG.gpu v0.9.1. Training data were subsetted with PLINK to contain only the SNPs present in the target data for CAAPA.

We limited the number of individuals within each reference panel to 200 to be able to compare the specific reference panels to the population reference panels. Indeed, this number is lower than the smallest population, allowing to resample the population and repeat the experiment.

### HLA imputation metrics

We have evaluated imputation accuracy using the F1-score. The F1-score is a harmonic mean of sensitivity (for a specific allele, # of correctly predicted allele/# of said alleles in the target dataset) and the positive predictive value (for a specific allele, # of correctly predicted allele/# of predictions of said allele). This score has the property to give important weight to the coverage of a specific allele prediction. For instance, if a rare allele is present once in a dataset of 100 alleles and not predicted by the model, you would have a 99% accuracy but a F1-score of 0.

*HLA* imputation models are limited by the pool of *HLA* alleles in the training dataset and the SNPs available, contrary to HLA-typing software based on read alignment, which relies on the complete database of known *HLA* alleles and the assessment of all gene regions. Therefore, we chose to average the results of all alleles present in the training and target datasets. Additionally, if one of these alleles is not predicted by the model, the positive predictive value, by definition, cannot be computed; in this case, the F1-score is also null. Since we wanted to focus our analysis on rare alleles, we decided to set the F1 scores of such alleles to 0, to visualize the impact of *HLA* alleles that are in the training dataset but do not manage to impute the ones in the target data.

### Dimension reduction

Principal Component Analysis (PCA) is routinely used in population genomics and association studies to study population ancestry. It relies on SNPs which are attributed to different contributions, maximizing the variance in their genotypes. It allows separating populations along multiple orthogonal axes with different contributions for each SNP. Uniform Manifold Approximation Projection (UMAP) and t-SNE are central in single-cell transcriptomics analyses (McInnes et al. 2018; Becht et al. 2018). Recently, It has also appeared in population genomics publications (Diaz-Papkovich et al. 2021; Sakaue et al. 2020). UMAP is based on simplicial topology to identify sets of neighbors for each individual and try to preserve them while transforming coordinates into new ones with less dimensions.

We performed dimension reduction after merging 1KG and CAAPA data. We ran PCA with PLINK, and UMAP on the BiRD cluster from Nantes University, using the umap R package. This package does not handle missing data; therefore, we applied the PLINK geno filter with a 0 threshold beforehand to remove any SNP with missing data. We followed the same process with SABE but merged the dataset with both 1KG and CAAPA.

We applied a silhouette score on the coordinates of the CAAPA individuals to identify the preferred number of clusters. We then performed k-means with the number of clusters that had the highest silhouette score. If the maximum score was inferior to 0.4, we chose not to perform clustering because simulations showed different groups would overlap greatly.

### Data access

1,000 Genomes SNP genotypes were retrieved from the International Genome Sample Resource (ISGR) and can be accessed through (https://www.internationalgenome.org/data-portal/data-collection/30x-grch38). 1,000 Genomes HLA genotypes of 2,693 individuals were recovered from Abi-Rached *et al*. (2018) at https://doi.org/10.1371/journal.pone.0206512.s010.

CAAPA SNPs were retrieved from the WGS data deposited in dbGAP with the accession code phs001123.v2.p1, described in Mathias, R. A. *et al*. (2016). CAAPA HLA genotypes were obtained with the Omixon software as described in https://doi.org/10.1016/j.jaci.2020.01.011.

For SABE, individual-level sequence datasets (BAM files) are available at the European Genome-phenome Archive (EGA), under EGA Study accession number EGAS00001005052. Further information about EGA can be found on https://ega-archive.org.

## Acknowledgements

We would like to thank the Centre de Calcul Intensif des Pays de la Loire. This work was supported by de Nantes Métropole, Région des Pays de la Loire and European Union (FEDER) via the Programme d’investissements d’Avenir (NExT, SHLARC Project, Nantes Université), the ATIP-Avenir Inserm program, the Région Pays de Loire ConnectTalent. Venceslas Douillard has received funding from the Inserm and Région Pays de la Loire. Nicolas Vince has received funding from the European Union’s Horizon 2020 research and innovation program under the Marie Skłodowska-Curie grant agreement No. 846520. Funding was provided by The São Paulo Research Foundation (FAPESP) grants and fellowships (CEPID 2013/08028-1, SABE 2014/50649-6, INCT 2014/50931-3, 2013/17084-2, 2017/19223-0, 2012/24731-1, 2018/15579-8, 2015/25020-0, 2020/02413-4) and Conselho Nacional de Desenvolvimento Científico e Tecnológico (CNPq INCT 465355/2014-5). The grant 2013/17084-2 supported creating and maintaining the hla-mapper software and the pipeline from HLA calling from NGS data.

