## Supplemental figures for "Optimal HLA imputation of admixed population with dimension reduction"

#### Supplementary Data

**Figure S1:** PCA and UMAP representations of a dataset containing both CAAPA and 1KG populations, according to the SNPs used (the chromosome 6 without MHC, or inside the MHC) and the number of dimensions selected (2 or 10). (A) Two dimensions of a PCA based on chromosome 6 SNPs without MHC; (B) Two-dimensional UMAP based on chromosome 6 SNPs without MHC; (C) Two first dimensions of a ten-dimensional UMAP based on MHC SNPs; (D) Two first dimensions of a ten-dimensional UMAP based on chromosome 6 SNPs without MHC.

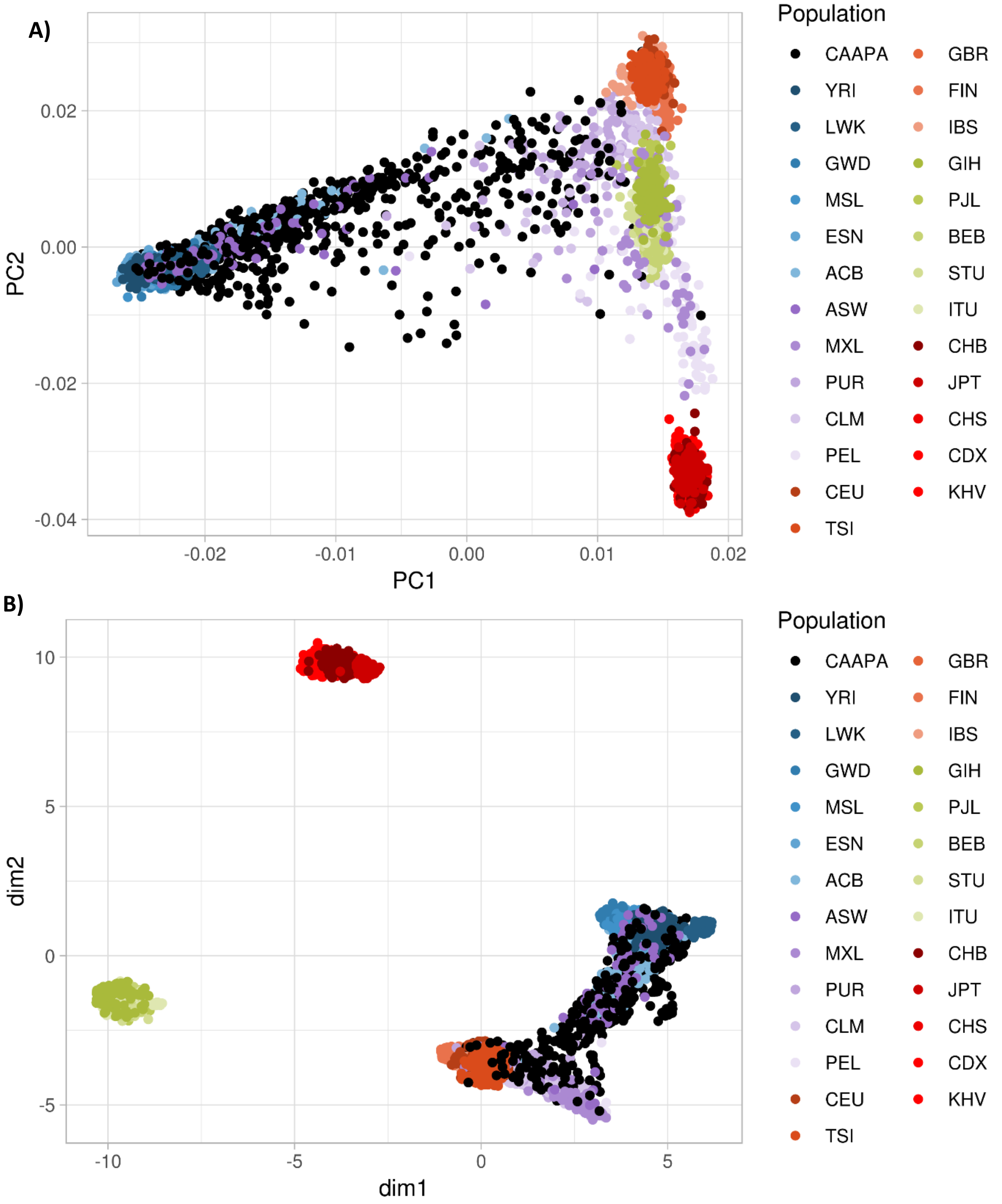

C)

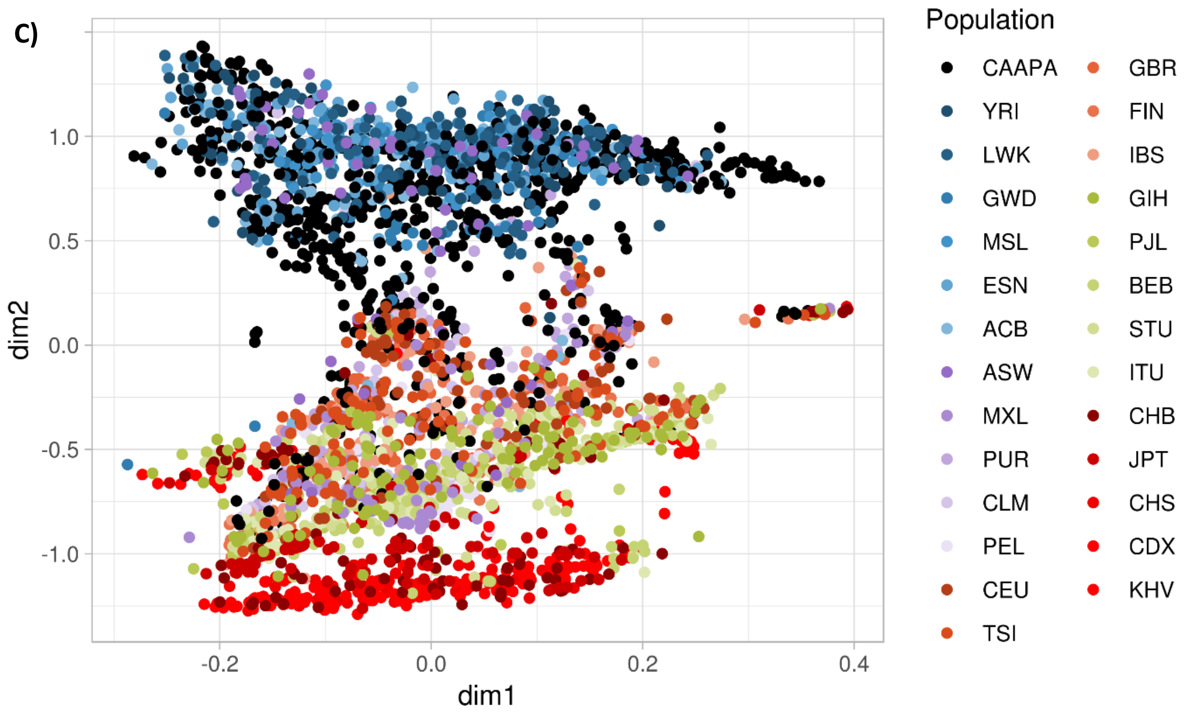
dim2
dim1
1.0
0.5
0.0
-0.5
-1.0
-0.2
0.0
0.2
0.4
Population
CAAPA
YRI
LWK
GWD
MSL
ESN
ACB
ASW
MXL
PUR
CLM
PEL
CEU
TSI
GBR
FIN
IBS
GIH
PJL
BEB
STU
ITU
CHB
JPT
CHS
CDX
KHV

D)

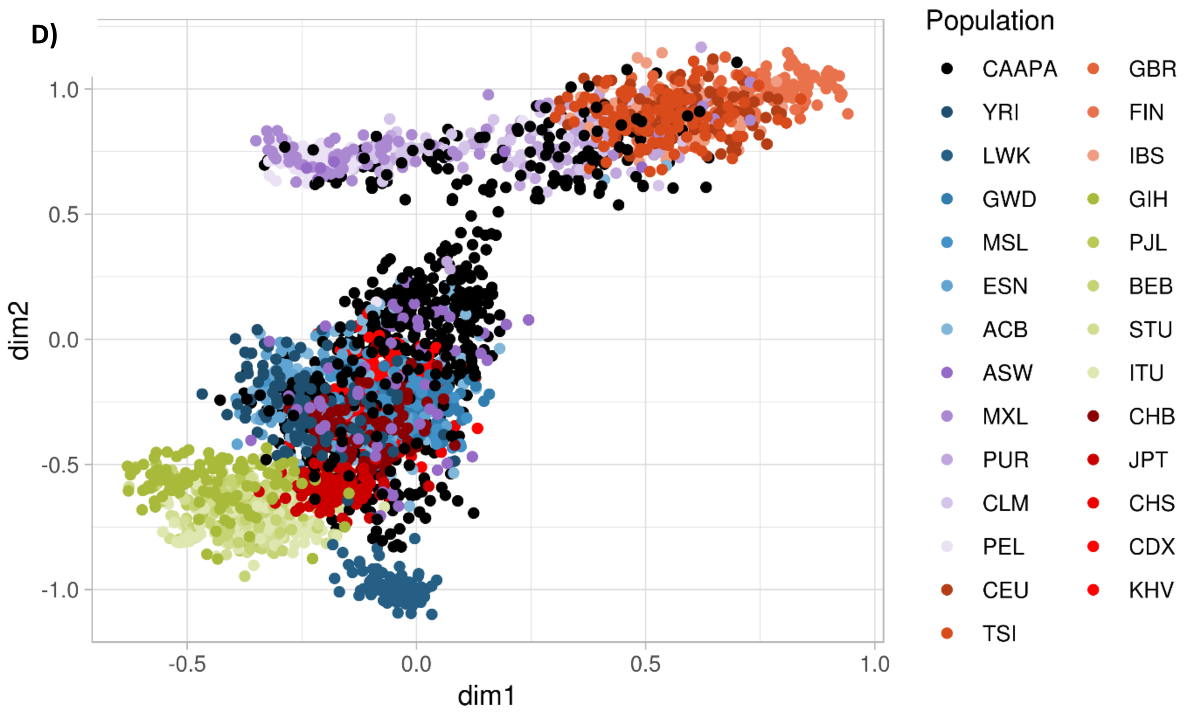
dim2
dim1
1.0
0.5
0.0
-0.5
-1.0
-0.5
0.0
0.5
1.0
Population
CAAPA
YRI
LWK
GWD
MSL
ESN
ACB
ASW
MXL
PUR
CLM
PEL
CEU
TSI
GBR
FIN
IBS
GIH
PJL
BEB
STU
ITU
CHB
JPT
CHS
CDX
KHV

**Figure S2:** Comparison of custom dataset composition, based on the sum of different individuals in the training dataset(s). Some datasets have more than 200 individuals because they represent multiple models of 200 individuals which were close from different subsets of CAAPA. Summing them up does not round to 400 or 600 because some individuals overlap. "Whole" datasets correspond to the closest 200 1KG individuals selected from averaging the positions of all CAAPA individuals in the PCA/UMAP representation, not from subsets. Nomenclature of the models can be found in table S2.

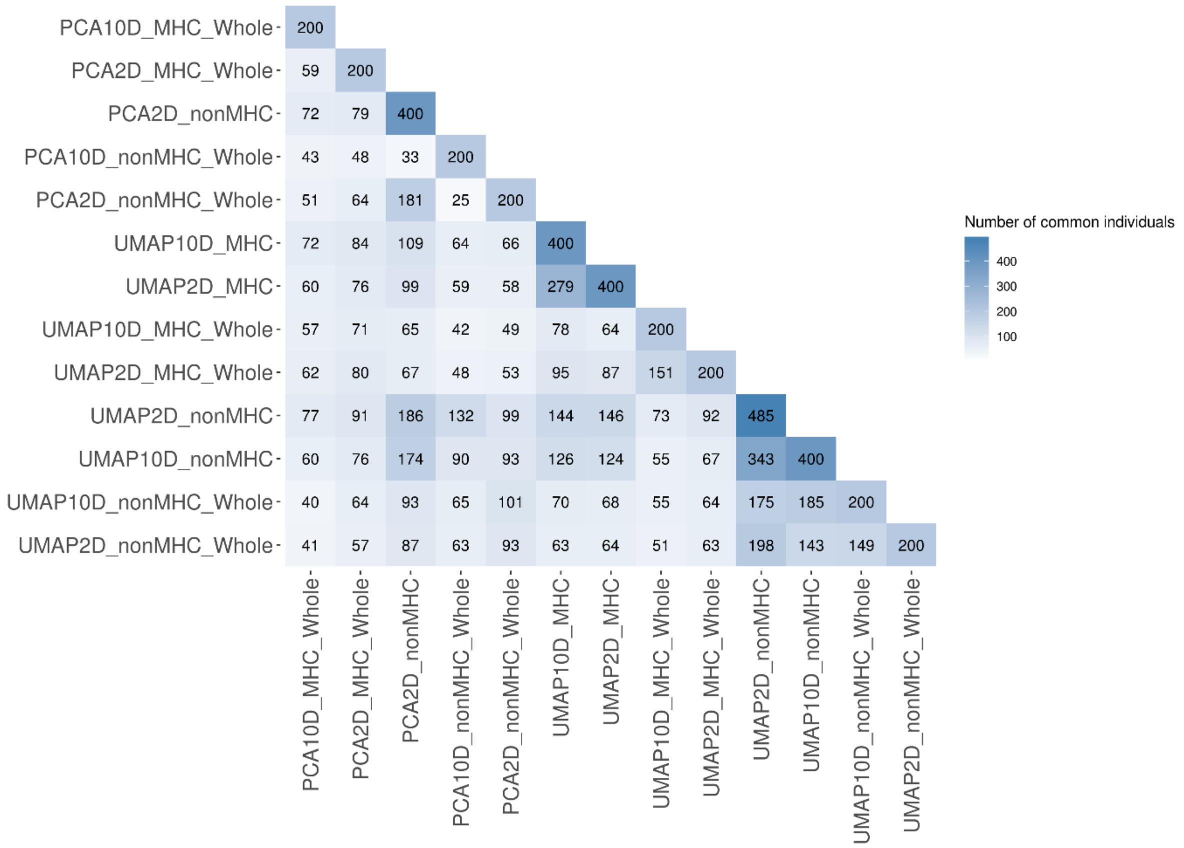

**Figure S3:** Accuracy measured by mean allelic error rate (1-Accuracy) for CAAPA HLA genotype imputation with the full 1KG, super-populations of 1KG, or the two custom models with the highest accuracy as training models. It represents the same information as accuracy but is easier to visualize: a higher value represents a lower HLA imputation accuracy. It is obtained by averaging each allele error rate.

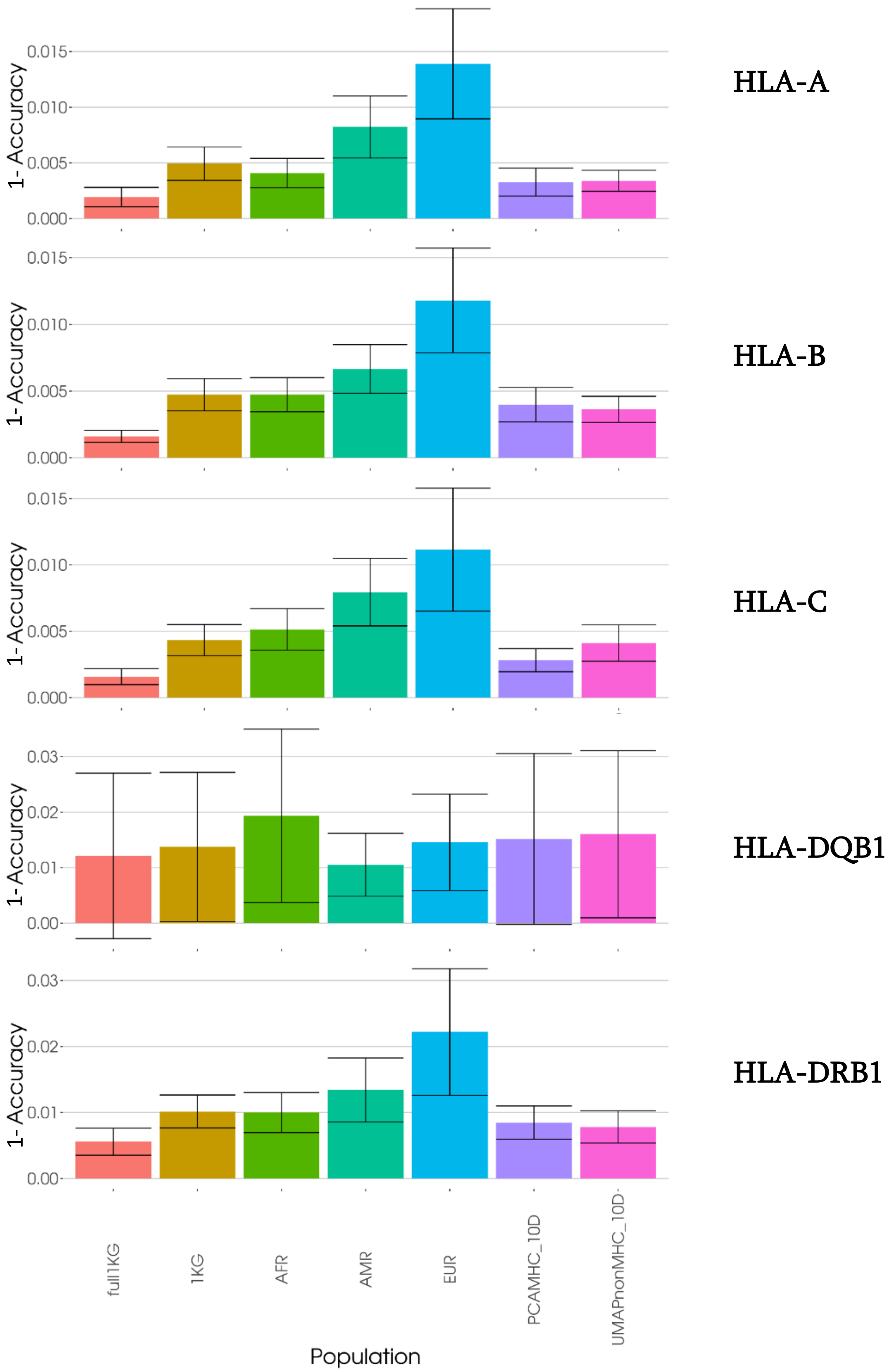

**Figure S4:** (A) Individual (counted as 0 or 1; incorrect vs. correct imputation) and (B) haplotype (individuals can be counted as 0, 0.5, or 1; incorrect vs. 1 correct allele vs. 2 correct alleles imputation) overall accuracies of CAAPA HLA genotypes predictions using 1KG as a training dataset.

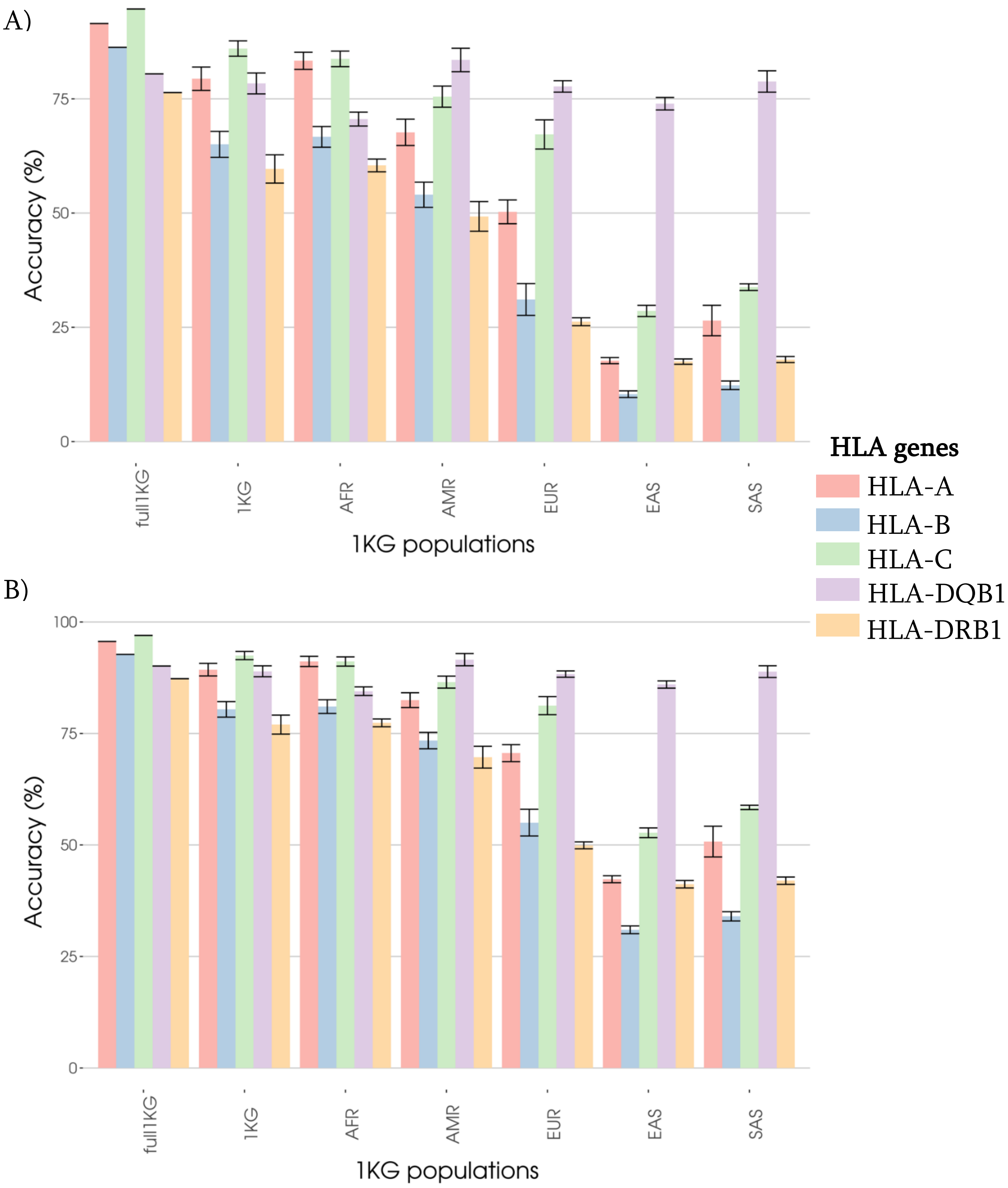

**Figure S5:** Mean F1-score by allele for each HLA gene *HLA-A*, *HLA-B*, *HLA-C*, *HLA-DQB1*, and *HLA-DRB1*. We compared the full 1KG dataset, to super-populations, to the most accurate custom models. HLA alleles that were absent from the 1KG dataset have been removed, therefore the alleles with no call at all mostly come from a singleton allele in the test dataset that has been mis-imputed once.

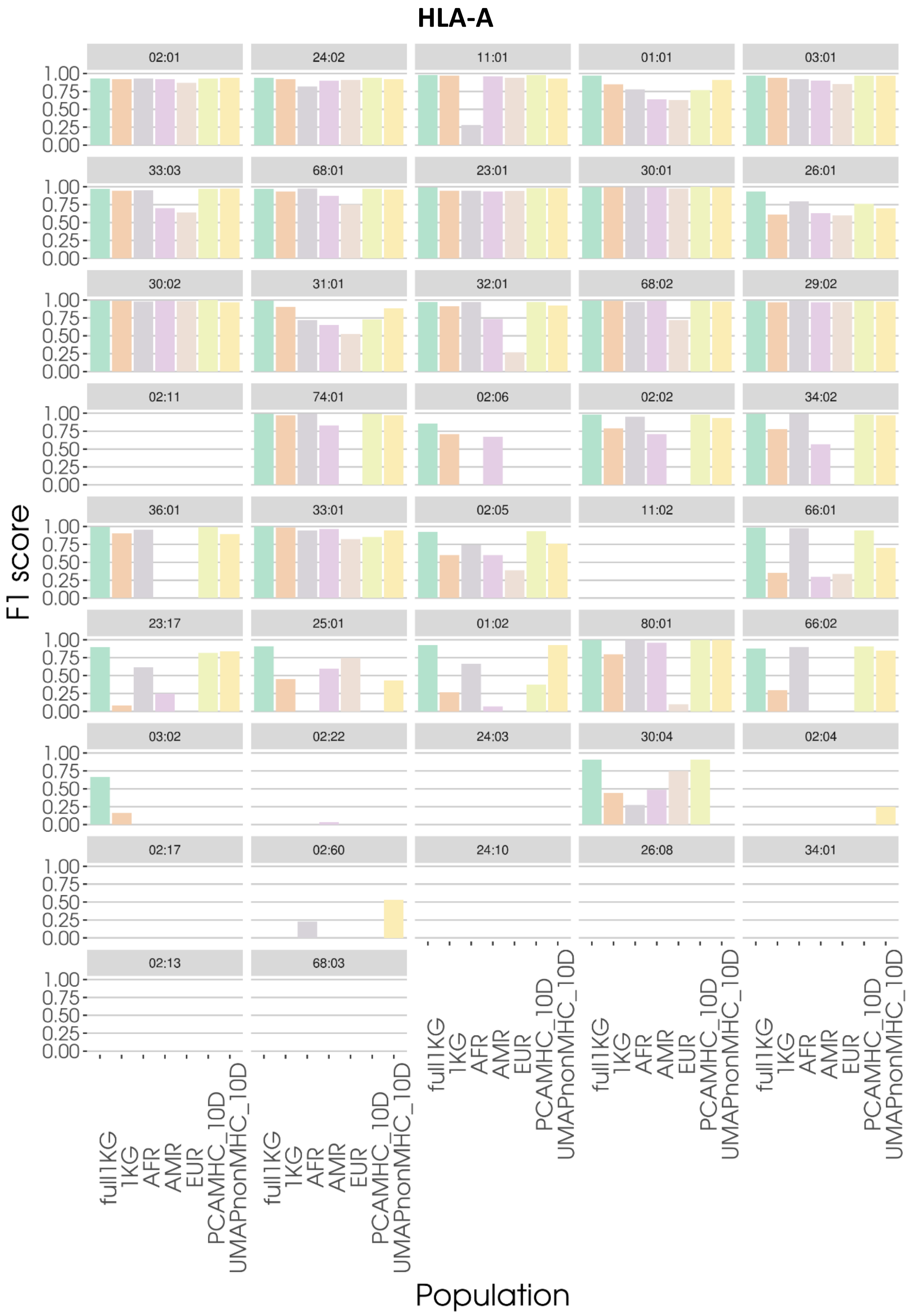

#### HLA-B

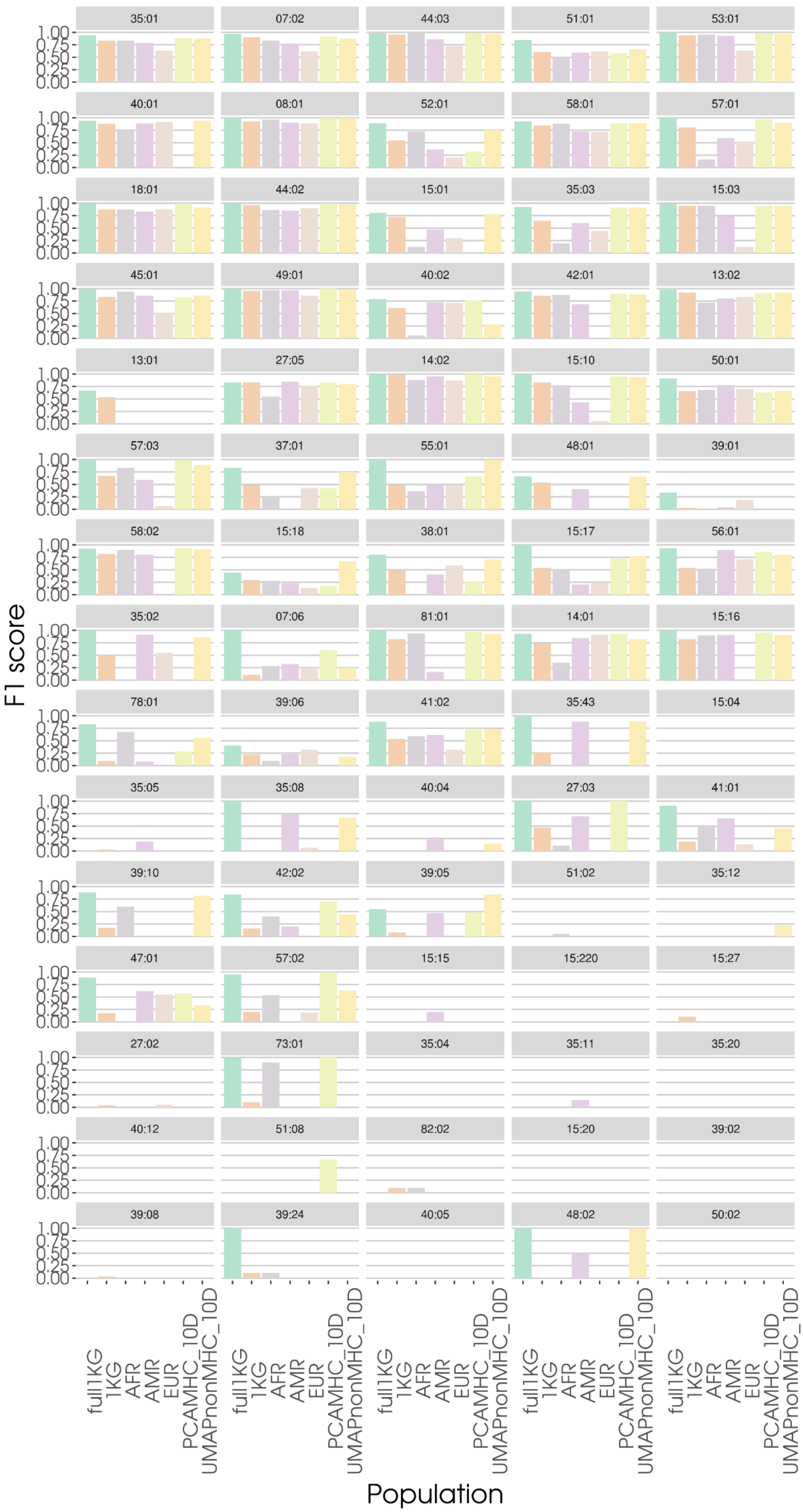
35:01
07:02
44:03
51:01
53:01
40:01
08:01
52:01
58:01
57:01
18:01
44:02
15:01
35:03
15:03
45:01
49:01
40:02
42:01
13:02
13:01
27:05
14:02
15:10
50:01
57:03
37:01
55:01
48:01
39:01
58:02
15:18
38:01
15:17
56:01
35:02
07:06
81:01
14:01
15:16
78:01
39:06
41:02
35:43
15:04
35:05
35:08
40:04
27:03
41:01
39:10
42:02
39:05
51:02
35:12
47:01
57:02
15:15
15:220
15:27
27:02
73:01
35:04
35:11
35:20
40:12
51:08
82:02
15:20
39:02
39:08
39:24
40:05
48:02
50:02
F1 score
1.00
0.75
0.50
0.25
0.00
full1KG
1KG
AFR
AMR
EUR
PCAMHC_10D
UMAPnonMHC_10D
Population

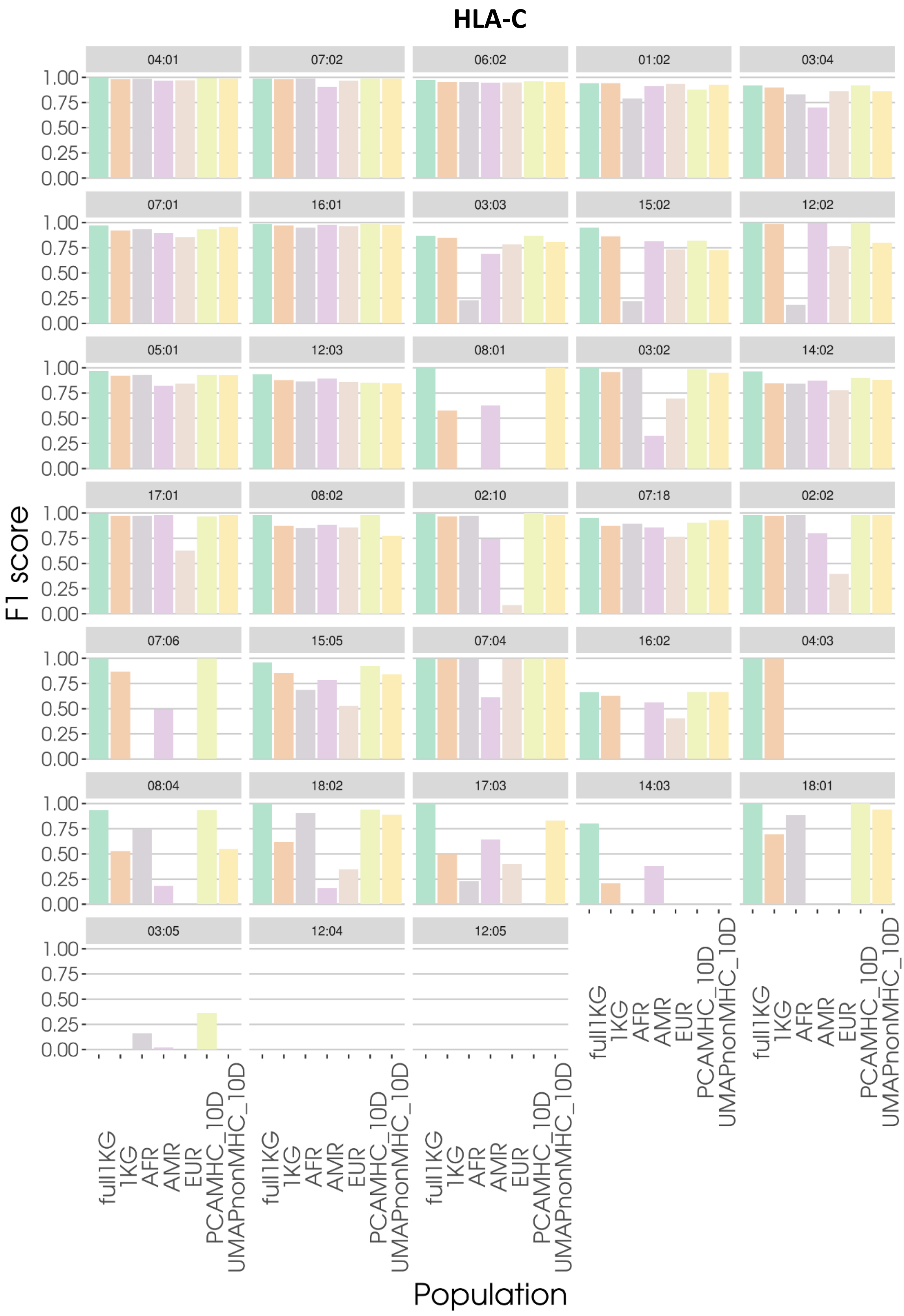
HLA-C
04:01
07:02
06:02
01:02
03:04
07:01
16:01
03:03
15:02
12:02
05:01
12:03
08:01
03:02
14:02
17:01
08:02
02:10
07:18
02:02
07:06
15:05
07:04
16:02
04:03
08:04
18:02
17:03
14:03
18:01
03:05
12:04
12:05
F1 score
1.00
0.75
0.50
0.25
0.00
full1KG
1KG
AFR
AMR
EUR
PCAMHC_10D
UMAPnonMHC_10D
Population

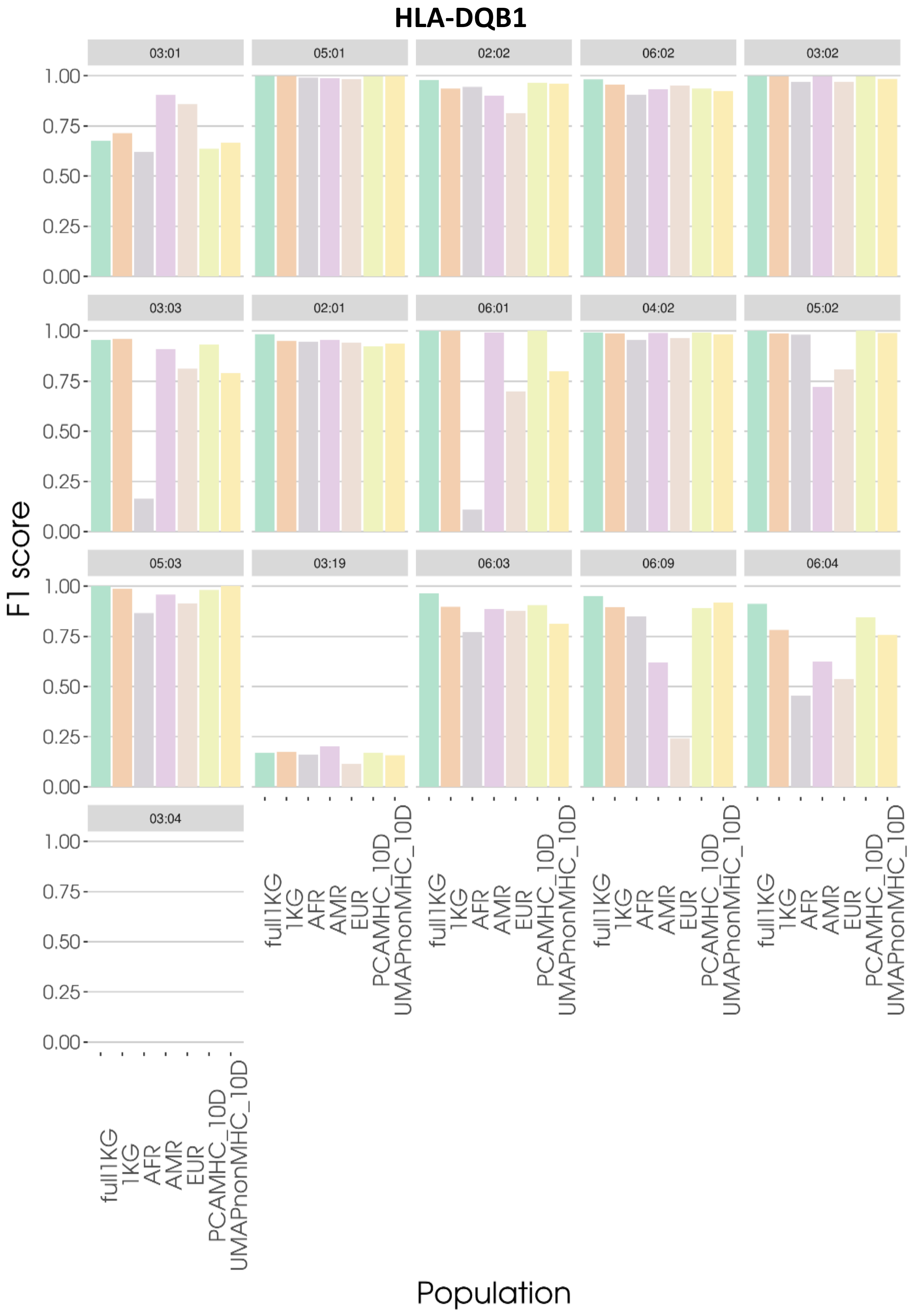

HLA-DQB1
03:01
05:01
02:02
06:02
03:02
03:03
02:01
06:01
04:02
05:02
05:03
03:19
06:03
06:09
06:04
03:04
1.00
0.75
0.50
0.25
0.00
F1 score
full1KG
1KG
AFR
AMR
EUR
PCAMHC_10D
UMAPnonMHC_10D
Population

### HLA-DRB1

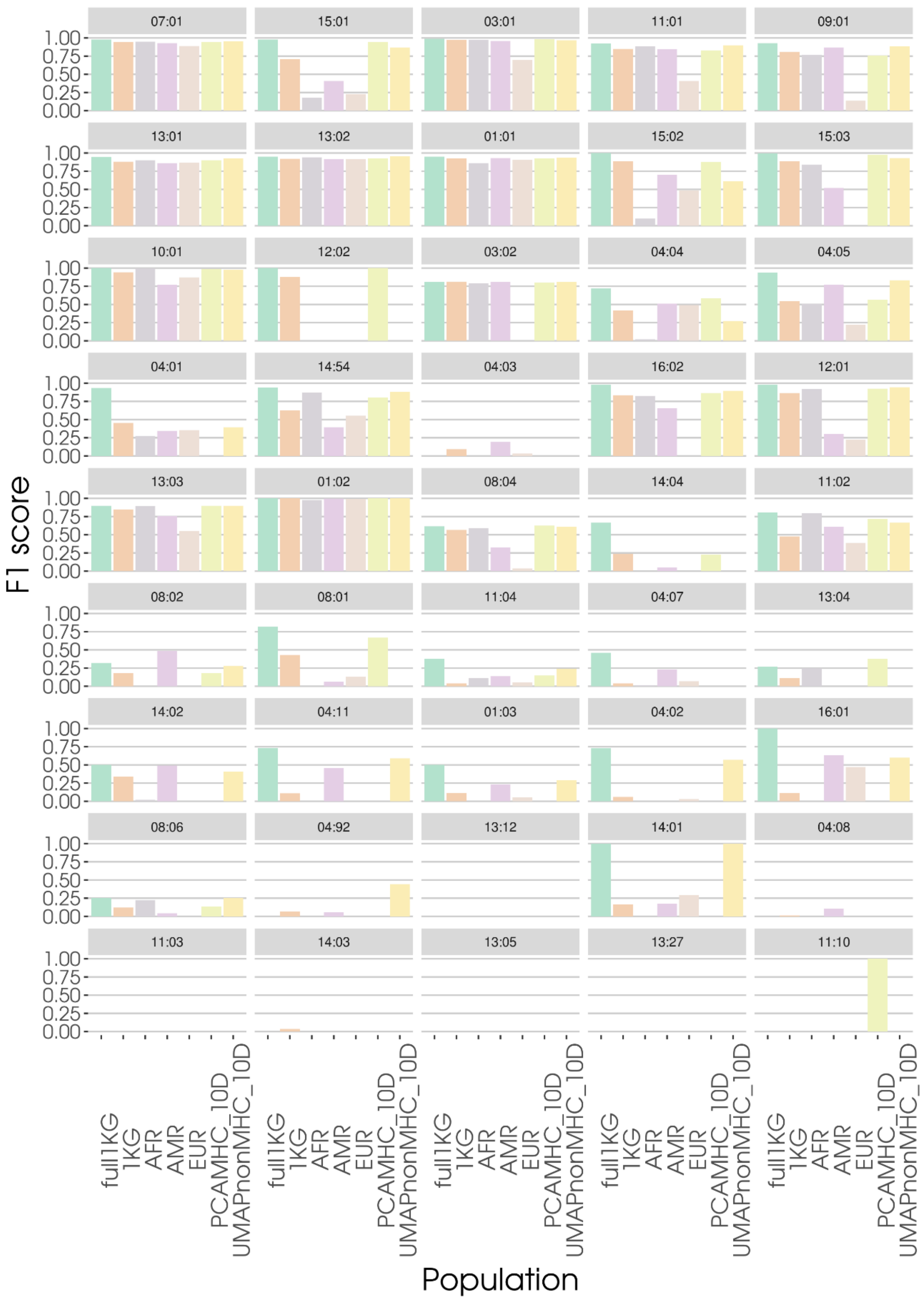

F1 score
Population
07:01
15:01
03:01
11:01
09:01
13:01
13:02
01:01
15:02
15:03
10:01
12:02
03:02
04:04
04:05
04:01
14:54
04:03
16:02
12:01
13:03
01:02
08:04
14:04
11:02
08:02
08:01
11:04
04:07
13:04
14:02
04:11
01:03
04:02
16:01
08:06
04:92
13:12
14:01
04:08
11:03
14:03
13:05
13:27
11:10
0.00
0.25
0.50
0.75
1.00
full1KG
1KG
AFR
AMR
EUR
PCAMHC_10D
UMAPnonMHC_10D

**Figure S6:** Global imputation accuracy of *HLA-B* and post-probability comparison between the full 1KG dataset and the one created with three models selected by ten-dimensional UMAP with SNPs outside of the MHC region. The x-axis shows the four cases where no model, one of them, or both, imputed individuals of CAAPA correctly. Percentages refer to the whole CAAPA dataset. Colors show for each case the number of genotypes which have a better post-probability either in the full or the custom 1KG model.

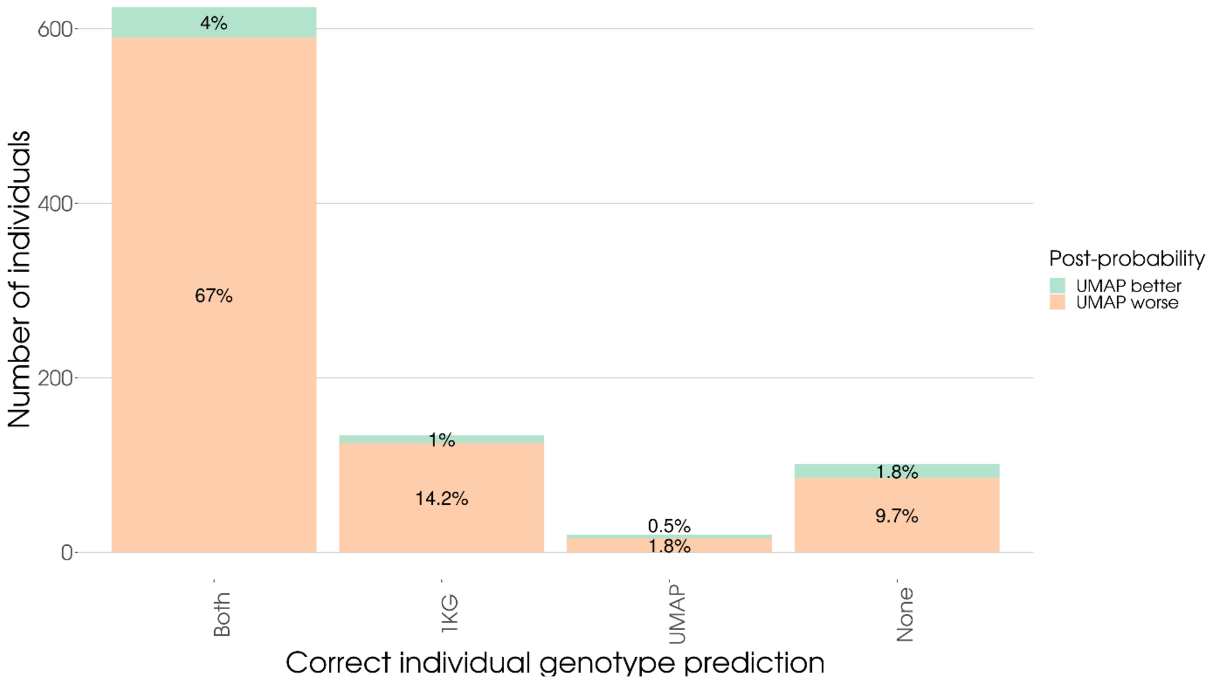

**Table S1**: Repartition of 1KG individuals in the 5 super-populations

| Super-population | Population |
|---|---|
| **AFR** - African ($n_{AFR}$=661) | **ACB** - African Carribean in Barbados ($n_{ACB}$=96) |
| | **ASW** - African ancestry in SouthWest US ($n_{ASW}$=61) |
| | **ESN** - Esan in Nigeria ($n_{ESN}$=99) |
| | **GWD** - Gambian in Western Division, The Gambia ($n_{GWD}$=113) |
| | **LWK** - Luhya in Webuye, Kenya ($n_{LWK}$=99) |
| | **MSL** - Mende in Sierra Leone ($n_{MSL}$=85) |
| | **YRI** - Yoruba in Ibadan, Nigeria ($n_{YRI}$=108) |
| **AMR** - American ($n_{AMR}$=347) | **CLM** - Colombian in Medellin, Colombia ($n_{CLM}$=94) |
| | **MXL** - Mexican Ancestry in Los Angeles, California ($n_{MXL}$=64) |
| | **PEL** - Peruvian in Lima, Peru ($n_{PEL}$=85) |
| | **PUR** - Puerto Rican in Puerto Rico ($n_{PUR}$=104) |
| **EAS** - East Asian ($n_{EAS}$=504) | **CDX** - Chinese Dai in Xishuangbanna, China ($n_{CDX}$=93) |
| | **CHB** - Han Chinese in Beijing, China ($n_{CHB}$=103) |
| | **CHS** - Southern Han Chinese, China ($n_{CHS}$=105) |
| | **JPT** - Japanese in Tokyo, Japan ($n_{JPT}$104) |
| | **KHV** - Kinh in Ho Chi Minh City, Vietnam ($n_{KHV}$=99) |
| **EUR** - European ($n_{EUR}$=503) | **CEU** - Utah residents with Norther and Western European ancestry ($n_{CEU}$=99) |
| | **FIN** - Finnish in Finland ($n_{FIN}$=99) |
| | **GBR** - British in England and Scotland ($n_{GBR}$=91) |
| | **IBS** - Iberian populations in Spain ($n_{IBS}$=107) |
| | **TSI** - Toscani in Italy ($n_{TSI}$=107) |
| **SAS** - South Asian ($n_{SAS}$=489) | **BEB** - Bengali in Bangladesh ($n_{BEB}$=86) |
| | **GIH** - Gujarati Indian in Houston, Texas ($n_{GIH}$=103) |
| | **ITU** - Indian Telugu in the UK ($n_{ITU}$=102) |
| | **PJL** - Punjabi in Lahore, Pakistan ($n_{PJL}$=96) |
| | **STU** - Sri Lankan Tamil in the UK ($n_{STU}$=102) |

**Table S2:** Nomenclature of the models. Two main dimension reduction methods have been applied to the SNP genotypes datasets, PCA and UMAP. They used the same set of SNP only taking the whole chromosome 6 without the MHC region, or the MHC region alone (29-34Mb) in the models. Then, we ran a k-means clustering method on two or ten dimensions of the reduced dataset to obtain subsamples of each model. For UMAPnonMHC_10D, we identified three clusters with a silhouette score on the ten dimensions, we ran the k-means on the dataset and created a sub-model for each of the groups (UMAPnonMHC_10D_1, UMAPnonMHC_10D_2 & UMAPnonMHC_10D_3). We then pooled the results to compute the accuracies of the whole model with other, non-custom, models. "Whole" datasets correspond to the closest 200 1KG individuals selected from averaging the positions of all CAAPA individuals in the PCA/UMAP representation, not from subsets.

| Name | Method | SNPs used | # of dimensions used for clustering | Custom or selected on the whole CAAPA dataset |
|---|---|---|---|---|
| UMAPnonMHC_2D | UMAP | Chromosome 6 without the MHC | 2 | Custom |
| UMAPnonMHC_10D | UMAP | Chromosome 6 without the MHC | 10 | Custom |
| UMAPMHC_2D | UMAP | Only the MHC region | 2 | Custom |
| UMAPMHC_10D | UMAP | Only the MHC region | 10 | Custom |
| PCAnonMHC_2D | PCA | Chromosome 6 without the MHC | 2 | Custom |
| PCAnonMHC_10D | PCA | Chromosome 6 without the MHC | 10 | Custom |
| PCAMHC_2D | PCA | Only the MHC region | 2 | Custom |
| PCAMHC_10D | PCA | Only the MHC region | 10 | Custom |
| UMAPnonMHC_whole2D | UMAP | Chromosome 6 without the MHC | 2 | Whole |
| UMAPnonMHC_ whole 10D | UMAP | Chromosome 6 without the MHC | 10 | Whole |
| UMAPMHC_ whole 2D | UMAP | Only the MHC region | 2 | Whole |
| UMAPMHC_ whole 10D | UMAP | Only the MHC region | 10 | Whole |
| PCAnonMHC_ whole 2D | PCA | Chromosome 6 without the MHC | 2 | Whole |
| PCAnonMHC_ whole 10D | PCA | Chromosome 6 without the MHC | 10 | Whole |
| PCAMHC_ whole 2D | PCA | Only the MHC region | 2 | Whole |
| PCAMHC_ whole 10D | PCA | Only the MHC region | 10 | Whole |
